# Revisiting the Fisher-KPP equation to interpret the spreading-extinction dichotomy

**DOI:** 10.1101/673202

**Authors:** Maud El-Hachem, Scott W. McCue, Wang Jin, Yihong Du, Matthew J. Simpson

## Abstract

The Fisher-KPP model supports travelling wave solutions that are successfully used to model numerous invasive phenomena with applications in biology, ecology, and combustion theory. However, there are certain phenomena that the Fisher-KPP model cannot replicate, such as the extinction of invasive populations. The Fisher-Stefan model is an adaptation of the Fisher-KPP model to include a moving boundary whose evolution is governed by a Stefan condition. The Fisher-Stefan model also supports travelling wave solutions; however, a key additional feature of the Fisher-Stefan model is that it is able to simulate population extinction, giving rise to a *spreading-extinction dichotomy.* In this work, we revisit travelling wave solutions of the Fisher-KPP model and show that these results provide new insight into travelling wave solutions of the Fisher-Stefan model and the spreading-extinction dichotomy. Using a combination of phase plane analysis, perturbation analysis and linearisation, we establish a concrete relationship between travelling wave solutions of the Fisher-Stefan model and often-neglected travelling wave solutions of the Fisher-KPP model. Furthermore, we give closed-form approximate expressions for the shape of the travelling wave solutions of the Fisher-Stefan model in the limit of slow travelling wave speeds, *c* ≪ 1.

## 1 Introduction

The Fisher-KPP model [1, 2, 3, 4, 5, 6] is a one-dimensional reaction-diffusion equation combining linear diffusion with a nonlinear logistic source term,

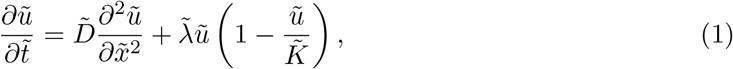

where 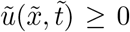 is the population density that depends on position 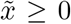, and time 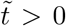. The dimensional parameters in Fisher-KPP model are the diffusivity 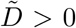, the proliferation rate 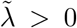 and the carrying capacity density 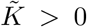. Solutions of the Fisher-KPP model on a semi-infinite domain that evolve from initial conditions with compact support asymptote to a travelling wave with a minimum wave speed, 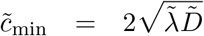 in the long time limit, 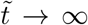 [1, 2, 3, 4, 5, 6]. The Fisher-KPP model also gives rise to travelling wave solutions with 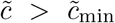 for initial conditions that decay sufficiently slowly as 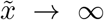 [1, 2, 3, 4, 5, 6], although for most practical applications we are interested in travelling wave solutions that move with the minimum wave speed since initial conditions with compact support are more often relevant [7, 8, 9, 10].

The Fisher-KPP model and its extensions have been used successfully in a wide range of applications including the study of spatial spreading of invasive species in ecology [11, 12, 13, 14]. In cell biology, the spatial spreading of invasive cell populations has been modelled using the Fisher-KPP model and its extensions for a range of applications including *in vitro* cell biology experiments [15, 16, 17, 18, 19] and in *vivo* malignant spreading [20, 21, 22]. Other areas of application include combustion theory [23, 24] and bushfire invasion [25]. Some of the extensions of the Fisher-KPP model involve working with different geometries [9, 16], such as inward and outward spreading in geometries with [26] and without [27] radial symmetry. Other variations include: (i) considering models with nonlinear diffusivity [28, 29, 30, 31, 32, 33]; (ii) incorporating different nonlinear transport mechanisms [34, 35, 36]; (iii) models of multiple invading subpopulations [31, 37]; and (iv) multi-dimensional models incorporating anisotropy [38]. The Fisher-KPP model gives rise to travelling wave-like solutions that do not allow the solution to go extinct. A cartoon of this kind of behaviour is shown schematically in Figure 1(a)-(c) where an invading cell population will propagate indefinitely on a semi-infinite domain.

**Figure 1:**
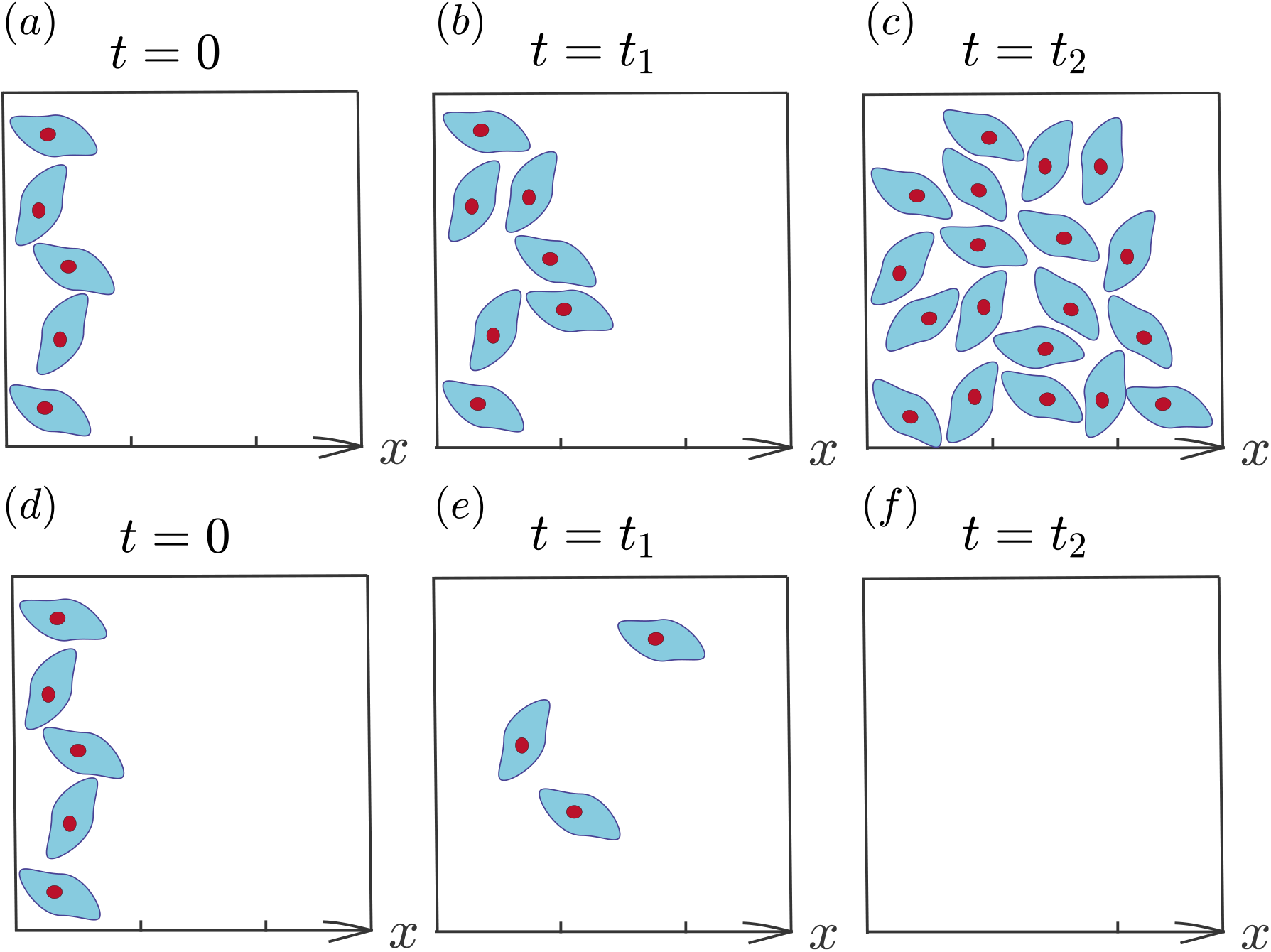
Schematic showing the evolution of a cell population. *(a)-(c)* The evolution of an invading cell population at *t* = 0, *t*_1_, and *t*_2_, with *t*_2_ > *t*_1_ > 0. In this case the population invades in the positive *x* direction indefinitely, provided that the domain is infinite. *(d)-(f)* The evolution of a cell population at *t* = 0, *t*_1_, and *t*_2_, with *t*_2_ > *t*_1_ > 0. In this case the population tends to extinction. Note that we have deliberately made the initial distribution of cells in (*a*) and (*d*) identical.

We are concerned here with the Fisher-Stefan model [39, 40, 41, 42, 43, 44, 45]

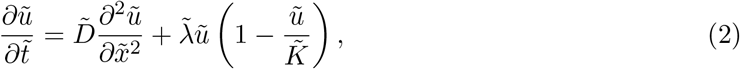

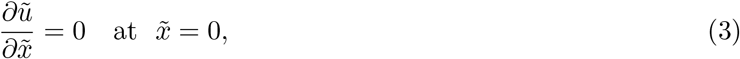

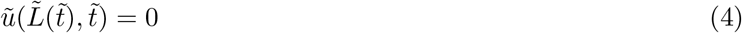

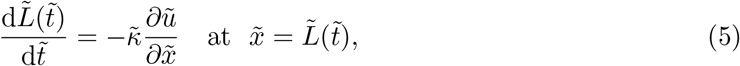

where 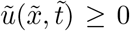 is the population density that depends on position 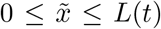 and time 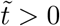. The parameters in Fisher-Stefan model are the same as in the Fisher-KPP model, as well as the Stefan parameter 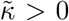, which relates the time rate of change of 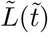 with the spatial gradient of the density, 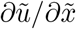, at the moving boundary 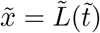.

As with the Fisher-KPP model, solutions of the Fisher-Stefan model, (2)–(5), can also lead to constant speed, constant shape travelling waves in the long time limit, as 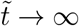 [39, 40, 41, 42, 43, 44, 45]. Interestingly, for the same initial condition but different choice of parameter ĩ, the Fisher-Stefan model can also give rise to a very different outcome whereby the population tends to extinction, 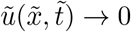 on 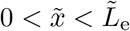 as 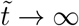 [39, 40, 41, 42, 43, 44, 45]. This major difference between the Fisher-KPP model and Fisher-Stefan model is of great interest because the Fisher-KPP model never leads to extinction, regardless of the choice of parameters. One way of interpreting this difference is that the Fisher-Stefan model is able to capture and predict additional details that are of practical interest because it is well known that many initially small translocated populations will become extinct [46]. This is one of the shortcomings of the Fisher-KPP model since this model implies that every small initial population always leads to successful invasion.

The Fisher-Stefan model is an adaptation of the Fisher-KPP model that includes a moving boundary, *x* = *L*(*t*), inspired by the classical Stefan problem [47, 48]. The classical Stefan problem is a one-dimensional model of heat conduction that includes a phase change, such as the conduction of heat associated with the melting of ice into water [47, 48]. An interesting mathematical and physical feature of the classical Stefan problem is that the interface between the two phases can move with time, giving rise to the notion of a *moving boundary problem* [47, 48]. Unlike classical models of heat conduction without any phase change [49], the solution of the Stefan problem requires the specification of two boundary conditions at the moving interface [47, 48]. First, the temperature at which the phase change occurs is specified at the moving boundary. This is analogous to Equation (3) in the Fisher-Stefan model. Second, the Stefan condition specifies a balance of latest heat energy to specific heat energy at the moving boundary, relating the time rate of change of position of the moving boundary to the flux of heat at the boundary [47, 48]. This is analogous to Equation (5) in the Fisher-Stefan model.

The moving boundary problem (2)–(5) with 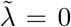 represents a one-phase Stefan problem which has an initial domain of solid, 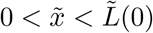, at some initial temperature 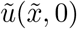, insulated at 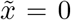. The interval 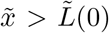 is initially occupied by liquid assumed to already be at the fusion temperature. For this particular problem formulation, the interface 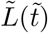 propagates into the liquid region as the heat energy contained within the solid is continually used as latent energy to convert the liquid to solid. The process continues until 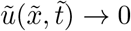 and 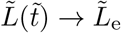 as 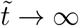, where a simple energy balance shows that

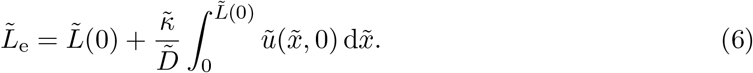

We shall return to this result later. A more general two-phase Stefan problem involves heat conduction in both phases, again separated by a moving boundary [47, 48]. Just like the Fisher-KPP model, there are many extensions of the classical Stefan problem such as dealing with higher-dimensions [50, 51, 52, 53] as well as modifying the moving boundary condition [54].

## 2 Results and discussion

### 2.1 Nondimensionalisation

We introduce the dimensionless variables, 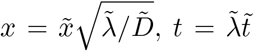 and 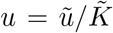, to rescale the Fisher-KPP equation as

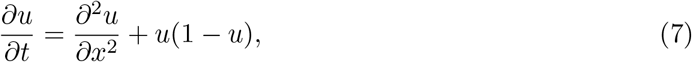

on *x* ≥ 0 and *t* > 0. It is useful to note that Equation (7) involves no free parameters, and that all solutions of Equation (7) with compactly supported initial conditions will eventually lead to travelling waves that move with speed *c*_min_ = 2. In this work we always specify the initial condition to be

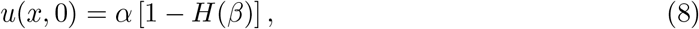

where *H*(*x*) is the usual Heaviside function, and *α* and *β* are positive constants so that we have *u*(*x*, 0) = *α* for *x* < *β* and *u*(*x*, 0) = 0 for *x* > *β*. All numerical solutions of Equation (7) that we consider specify *∂u/∂x* = 0 at *x* = 0 and *∂u/∂x* = 0 at *x* = *x*_∞_. Here *x*_∞_ is chosen to be sufficiently large so that we can numerically approximate the infinite domain problem on 0 ≤ *x* < ∞ by the finite domain problem 0 ≤ *x* ≤ *x*_∞_ [31]. Full details of the numerical method used to solve Equation (7), together with benchmark test cases to confirm the accuracy of our numerical solutions are given in the Supplementary Material.

To nondimensionalise the Fisher-Stefan model we employ the same dimensionless variables as in the Fisher-KPP model with 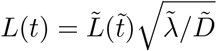 and 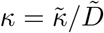 so that we have

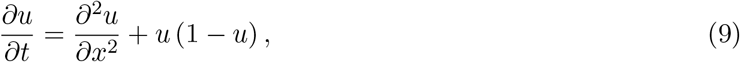

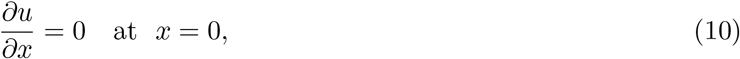

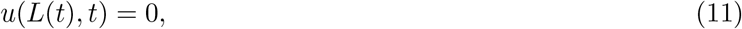

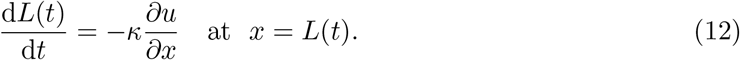

It is relevant to note that the nondimensional Fisher-Stefan model involves one parameter, *κ* > 0. For all numerical solutions of Equations (9)–(12) in this work we always apply the initial condition (8) such that *L*(0) = *β*. Full details of the numerical method used to solve Equations (9)–(12), together with benchmark test cases to confirm the accuracy of our numerical solutions, are given in the Supplementary Material.

To illustrate key features of the Fisher-KPP and Fisher-Stefan models we present numerical solutions of both models in Figure 2. Results in Figure 2(a) show the time evolution of the solution of the Fisher-KPP model from an initial condition with compact support. Plotting solutions at equally spaced values of time suggests that the solution approaches a travelling wave that moves in the positive *x* direction with constant speed and constant shape. Our numerical solutions confirm that the speed of propagation is *c* = 2, as expected. Results in Figure 2(b) show the time evolution of the solution of the Fisher-Stefan model (9)–(12) for the same initial condition used in Figure 2(a) together with a particular choice of *κ*. Again, plotting solutions at equally spaced values of time suggests that the solution approaches a travelling wave that moves in the positive *x* direction with constant speed and constant shape. In this case, for our choice of *κ*, our numerical solution suggests that *c* = 1.2, which is slower than the minimum wave speed for the Fisher-KPP model. Another important difference between the travelling wave solutions in Figure 2(a)-(b) is that the travelling wave solution of the Fisher-KPP model does not have compact support since *u*(*x, t*) > 0 for all *x* ≥ 0 and *t* > 0. This feature of the Fisher-KPP model has been previously criticised as being biologically implausible [7, 8, 30] (and this observation has motivated extensions of the Fisher-KPP model to include various nonlinear diffusion terms so that the resulting travelling waves have compact support [7, 8, 30]). In contrast, owing to the boundary conditions at *x* = *L*(*t*), the travelling wave solution of the Fisher-Stefan has compact support since we have *u*(*L*(*t*),*t*) = 0 for *t* > 0. Therefore, we have identified two features of the Fisher-Stefan model that are appealing when compared to the Fisher-KPP model: (i) the Fisher-Stefan model permits population extinction whereas the Fisher-KPP model implies that all initial populations successfully invade; and (ii) travelling wave solutions of the Fisher-Stefan model have compact support whereas travelling wave solutions of the Fisher-KPP model do not.

Results in Figure 2(c) show the time evolution of the solution of the Fisher-Stefan model for the same initial condition used in Figure 2(a)-(b), but this time we choose a slightly smaller value of *κ*. In this instance, plotting the solutions at the same intervals of time as in Figures 2(a)-(b) indicates that the solution does not tend towards a travelling wave, and instead appears to go extinct. Figure 2(d) shows a magnified view of the solution in Figure 2(c) so that we can see additional details as the population tends to extinction. From these magnified solutions we see that *L*(0) = 1. By *t* = 20 we have *L*(20) ≈ 1.4, and after this time the solution rapidly tends to zero. Together, the results in Figure 2(b)-(d) illustrate the spreading-extinction dichotomy since for certain choices of *κ* we observe spreading as a travelling wave in Figure 2(b), whereas keeping everything else identical except for choosing a smaller value of *κ* we observe the population going extinct in Figure 2(c)-(d). These initial comparisons in Figure 2 indicate that the Fisher-KPP and Fisher-Stefan models appear to be very different when they are described in terms of partial differential equations. Since both the Fisher-KPP and Fisher-Stefan models support travelling wave solutions we will now explore these models in the phase plane.

### 2.2 Phase plane analysis

Numerical solutions of the Fisher-KPP model in Figure 2(a) suggest that we seek travelling wave solutions with travelling wave coordinate *z* = *x* — *ct*, where *c* > 0 is the constant speed of propagation in the positive *x* direction. In the travelling wave coordinate, Equation (7) simplifies to a second order nonlinear ordinary differential equation

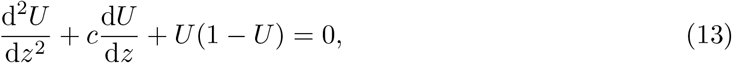

where –∞ < *z* < ∞, with *U*(–∞) = 1 and *U*(∞) = 0.

**Figure 2:**
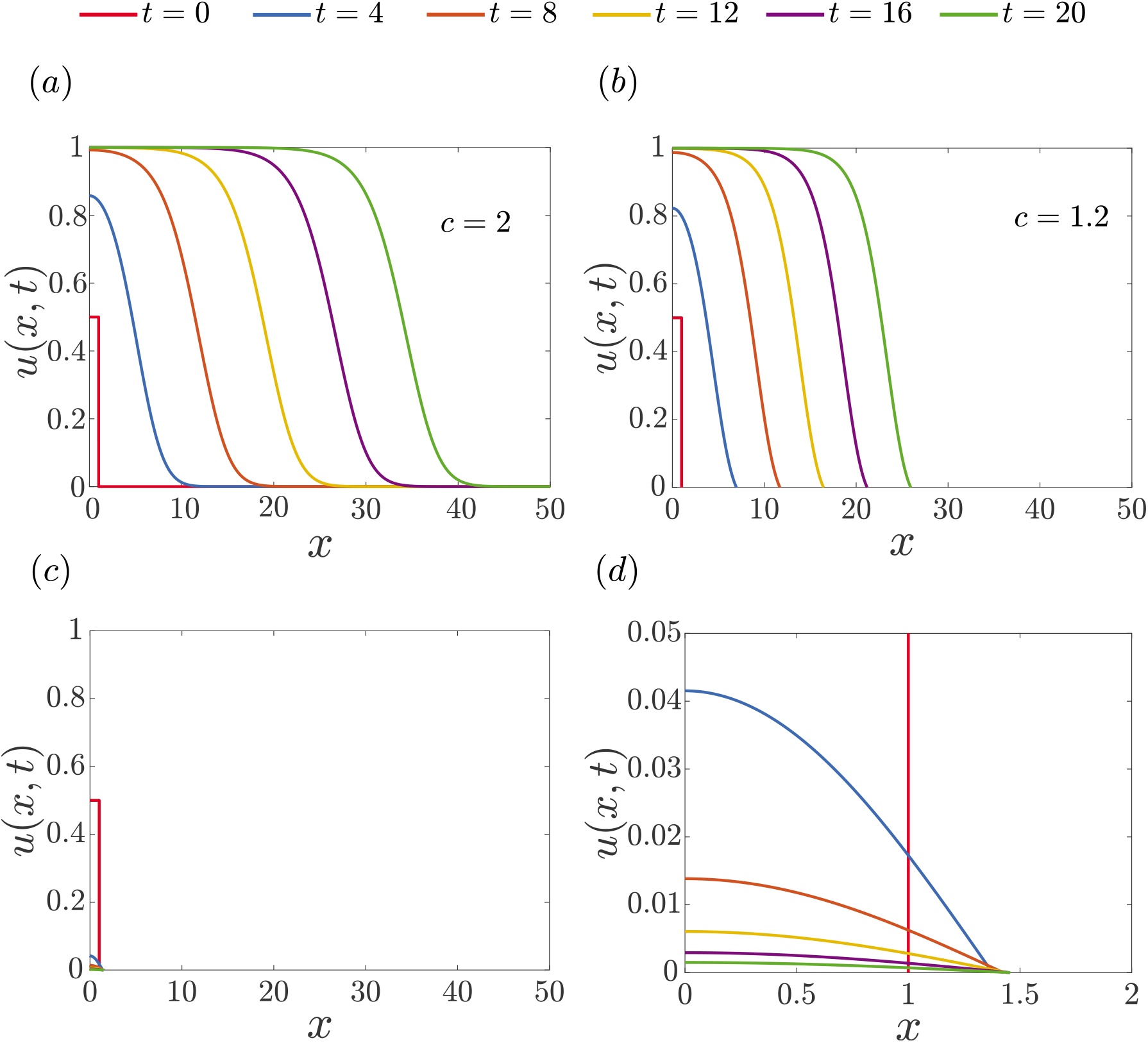
Numerical solutions of the Fisher-KPP and the Fisher-Stefan models. *(a)* Numerical solutions of the Fisher-KPP equation evolving into a travelling wave solution with the minimum wave speed, *c* = 2. *(b)* Numerical solutions of Fisher-Stefan model evolving into a travelling wave solution with *c* = 1.2. *(c)* Numerical solutions of the Fisher-Stefan model leading to extinction. *(d)* Magnified region of the solution in *(c)*, for 0 ≤ *x* ≤ 2, to clearly show the dynamics of the extinction behaviour. For the Fisher-Stefan model we set *κ* = 20 in *(b)* and *κ* = 0.45 in *(c).* Numerical solutions of the Fisher-Stefan model are obtained with Δ*ξ* = 1 × 10^-4^, whereas numerical solutions of the Fisher-KPP model are obtained with Δ*x* = 1 × 10^-4^. For both the Fisher-KPP and Fisher-Stefan models we set Δ*t* = 1 × 10^-3^ and *ϵ* = 1 × 10^-8^. For all results presented here the initial condition is Equation (8) with *α* = 0.5 and *β* = 1.

Our treatment of the analysis of travelling wave solutions of the Fisher-Stefan model is very similar except that we must first assume that our choice of initial condition and *κ* in Equation (9)–(12) is such that a travelling wave solution forms, as in Figure 2(b). In this case, writing Equation (9) in the travelling wave coordinate gives rise to the same second order nonlinear ordinary differential equation, Equation (13) with *U*(–∞) = 1. The other boundary condition is treated differently and to see this difference we express the Stefan condition (12) in terms of *z*, giving

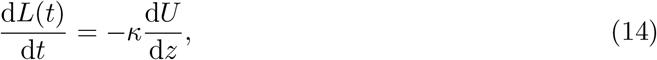

at *z* = *L*(*t*) – *ct*. For a travelling wave solution we have *d*L(*t*)/d*t* = *c*, so that the differential equation (13) holds on –∞ < *z* < 0. The boundary conditions are given by *U*(–∞) = 1, and

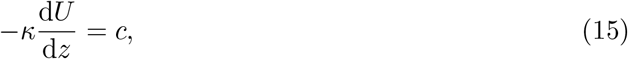

at *z* = 0. Therefore, while the Fisher-KPP and Fisher-Stefan models presented as partial differential equations appear to be very different, when we seek travelling wave solutions of these models in the travelling wave coordinate we find that the equations governing the phase planes for the two models are the same, with differences only at one boundary condition.

We first examine travelling wave solutions of the Fisher-KPP model by re-writing Equation (13) as a first order dynamical system

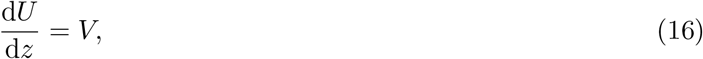

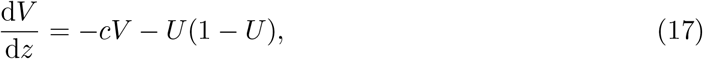

for –∞ < *z* < ∞. This dynamical system has two equilibrium points: (1, 0) and (0, 0). A travelling wave solution corresponds to a heteroclinic trajectory between these two equilibrium points. Linear analysis shows that the eigenvalues at (1,0) are 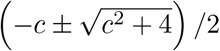 so that the local behaviour at (1, 0) is a saddle point [5]. The eigenvalues at (0,0) are 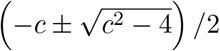, meaning that the local behaviour at (0, 0) is a stable spiral for *c* < 2 and a stable node if *c* ≥ 2 [5]. Therefore, to avoid nonphysical negative solutions near (0, 0) we require *c* ≥ 2, giving rise to the well known minimum wave speed for the Fisher-KPP model [5, 55].

The shape of the heteroclinic orbit between (0, 0) and (1, 0) are given by the solution of

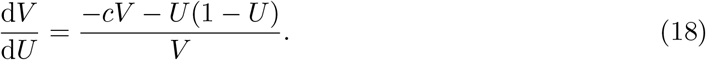

Neither the system (16)–(17) or Equation (18) have exact solutions for an arbitrary choice of *c* > 0. Therefore, we will consider numerical solutions of these ordinary differential equations when we present and visualise the phase planes in this work, with details of the numerical methods given in the Supplementary Material. The solution of Equation (18) gives *V*(*U*), while the solution of the system (16)-(17) gives *U*(*z*) and *V*(*z*), and we will use both approaches where relevant. All phase planes presented in this work are generated using a combination of exact and numerical methods that are outlined in the Supplementary Material.

Results in Figure 3 show a suite of phase planes for the Fisher-KPP model for a range of c. The phase plane in Figure 3(a) shows the flow, the location of the equilibrium points and the heteroclinic orbit for *c* = 10, which approaches (0, 0) without spiralling. When we plot the solution in terms of the density, *U* = *U*(*z*), in Figure 3(b), we see that this solution is positive and monotonically decreasing. Similar results are presented in Figure 3(c)-(d) for *c* = 2. In contrast, Figure 3(e) shows the phase plane for *c* = 0.5 where we see that the heteroclinic orbit approaches (0,0) as a spiral, indicating that *U*(*z*) < 0 in certain regions. This oscillatory behaviour is often invoked to justify the condition that *c* ≥ *c*_min_ = 2 for the Fisher-KPP model and the possibility of travelling waves with *c* < 2 is typically ignored [5].

Since travelling wave solutions of the Fisher-KPP and Fisher-Stefan models are governed by the same phase planes, it is worthwhile to examine how the phase planes in Figure 3 relate to the travelling wave solutions of the Fisher-Stefan model. As previously stated, travelling wave solutions of the Fisher-Stefan model satisfy a different boundary condition, (15). The trajectory in the phase plane must intersect with, and terminate at, (0, –*c*/*κ*). The phase planes and heteroclinic orbits for the Fisher-KPP model in Figure 3(a)-(b) and Figure 3(c)-(d) show that such an intersection is impossible for these choices of *c* ≥ 2. In particular, the linearisation about (0, 0) means that whenever *c* ≥ 2, the stable node at (0, 0) precludes the possibility of such an intersection, illustrating that there is no travelling wave solution for the Fisher-Stefan model with *c* ≥ 2. In contrast, for *c* < 2, the trajectory intersects the *V*-axis at some point, (0, –*c*/*κ*), as indicated by the orange disc in Figure 3(e). Therefore, this additional boundary condition for the Fisher-Stefan model together with the linearisation about (0, 0) indicates that travelling wave solutions for the Fisher-Stefan model require *c* < 2. Under these conditions, the trajectory in the phase plane corresponding to the travelling wave solutions runs between (1, 0) and (0, −*c/κ*), and since this trajectory never leaves the fourth quadrant we avoid the issue of negative densities. This is a very interesting result because the standard phase plane analysis of travelling wave solutions of the Fisher-KPP model typically discard any solutions for which *c* < 2 based on physical grounds [5, 4]. In this work we show that it is precisely these normally-discarded solutions that actually form the basis of the travelling wave solutions of the Fisher-Stefan model provided that we only consider that part of the trajectory between (1, 0) and (0, −*c/κ*), where *U*(*z*) ≥ 0. Therefore, by revisiting the travelling wave solutions of the Fisher-KPP model we are providing insight into the properties of travelling wave solutions of the Fisher-Stefan model.

**Figure 3:**
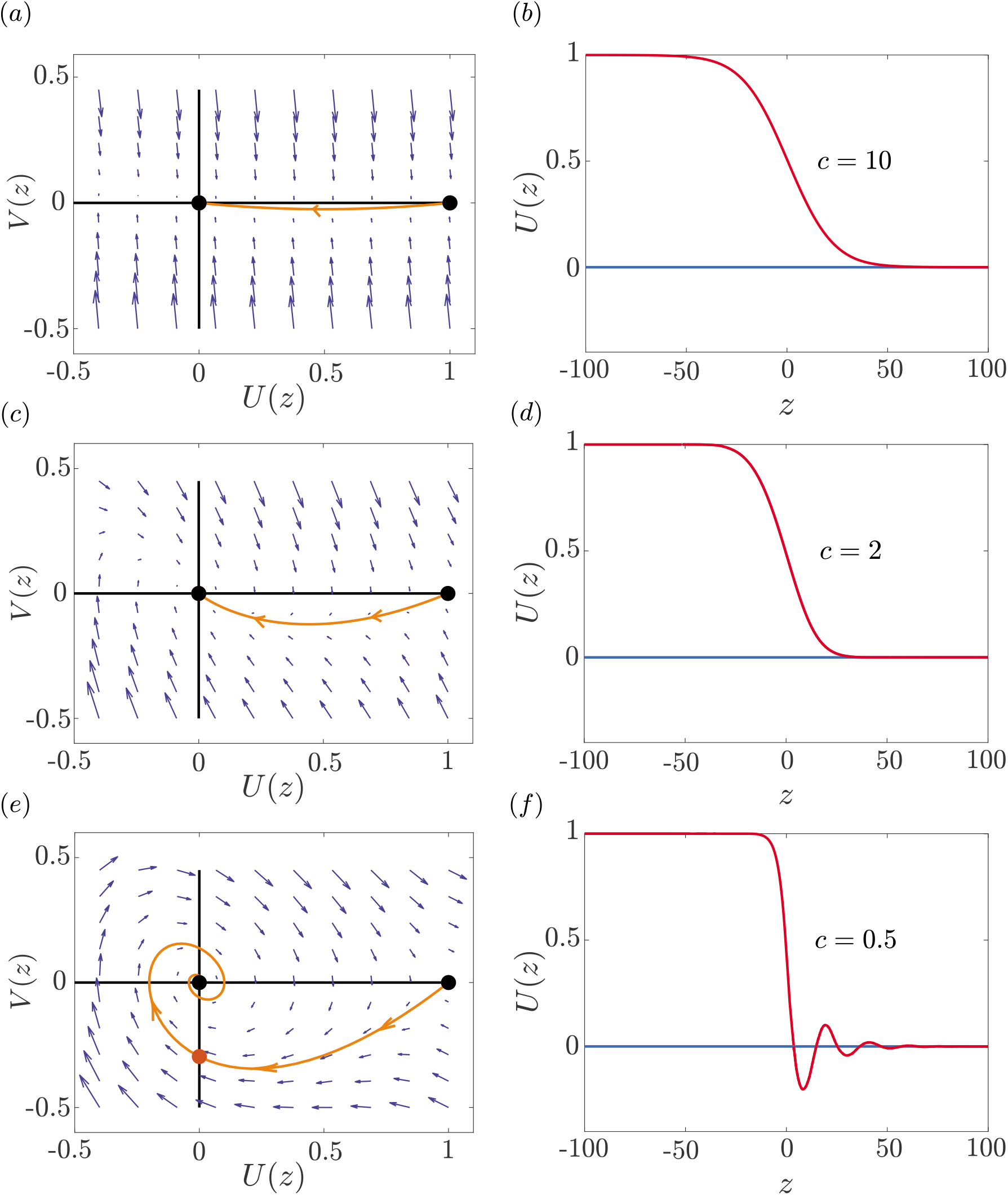
Phase planes and density profiles for the Fisher-KPP equation with various choices of *c*. *(a)* Phase plane and heteroclinic trajectory for *c* = 10. *(b)* The corresponding density profile for the heteroclinic trajectory in *(a*). *(c)* Phase plane and heteroclinic trajectory for *c* = 2. *(d)* The corresponding density profile for the heteroclinic trajectory in *(c). (e)* Phase plane and heteroclinic trajectory for *c* = 0.5. *(f)* The corresponding density profile for the heteroclinic trajectory in *(e)*. Equilibrium points at (1,0) and (0,0) are shown with black discs. The blue arrows show the flow associated with the dynamical system, and the solid orange line shows the heteroclinic trajectory that runs between (0, 0) and (1, 0). The orange disc in (c) shows the location where the heteroclinic trajectory intersects with the *U*(*z*) = 0 axis where *V*(*z*) < 0.

We now provide additional results in Figure 4 comparing trajectories in the phase plane for a wider range of *c*. The trajectories for *c* =10 and *c* = 2 show a heteroclinic orbit that runs between (1,0) and (0,0) without spiralling around the origin. These trajectories are associated with travelling wave solutions of the Fisher-KPP model for these choices of *c*. Additional results for *c* = 1 and *c* = 0.5 are also shown, and these trajectories clearly spiral near to the origin. However, both of these trajectories cross the *V*-axis at some point, shown with an appropriately coloured disc in Figure 4, where *U*(*z*) = 0 and *V*(*z*) < 0, which satisfies the Fisher-Stefan boundary condition. Therefore, the trajectories in Figure 4 with *c* < 2 are shown as a combination of solid and dashed lines. Those parts of the trajectories shown in solid correspond to the travelling wave solution of the Fisher-Stefan model, whereas the dashed parts of the trajectories are not associated with the travelling wave solution. Finally, we also include a trajectory in Figure 4 for *c* = 0. In this case the trajectory forms a homoclinic orbit with (1, 0). Although this trajectory does not correspond to any travelling waves with *c* > 0, later we will show it is important when constructing approximate perturbation solutions for *c* ≪ 1.

### 2.3 Relationship between *κ* and *c*

It is interesting to recall that all solutions of the Fisher-KPP model with compact support will always eventually form a travelling wave with the minimum, *c* = *c*_min_ = 2. In contrast, the question of whether travelling wave solution will form for the Fisher-Stefan model for a particular initial condition, and if so, what speed those travelling wave solutions move depends upon the choice of *κ*. To explore this relationship we use a combination of phase plane analysis and numerical solutions of the Fisher-Stefan model (9)–(12). By repeatedly solving the phase plane equations, (16) for different values of *c* < 2, we are able to estimate the point at which the trajectory leaving (1, 0) first intersects the *V*-axis and then use the boundary condition, *V* = −*c/κ* at *U* = 0, to calculate the corresponding values of *κ*. The solid blue curve in Figure 5 shows this relationship. We find that as we examine increasing values of *c* towards the threshold value of *c* = 2, *κ* appears to grow without bound. This numerical result suggests that *κ* → ∞ as *c* → 2^−^.

**Figure 4:**
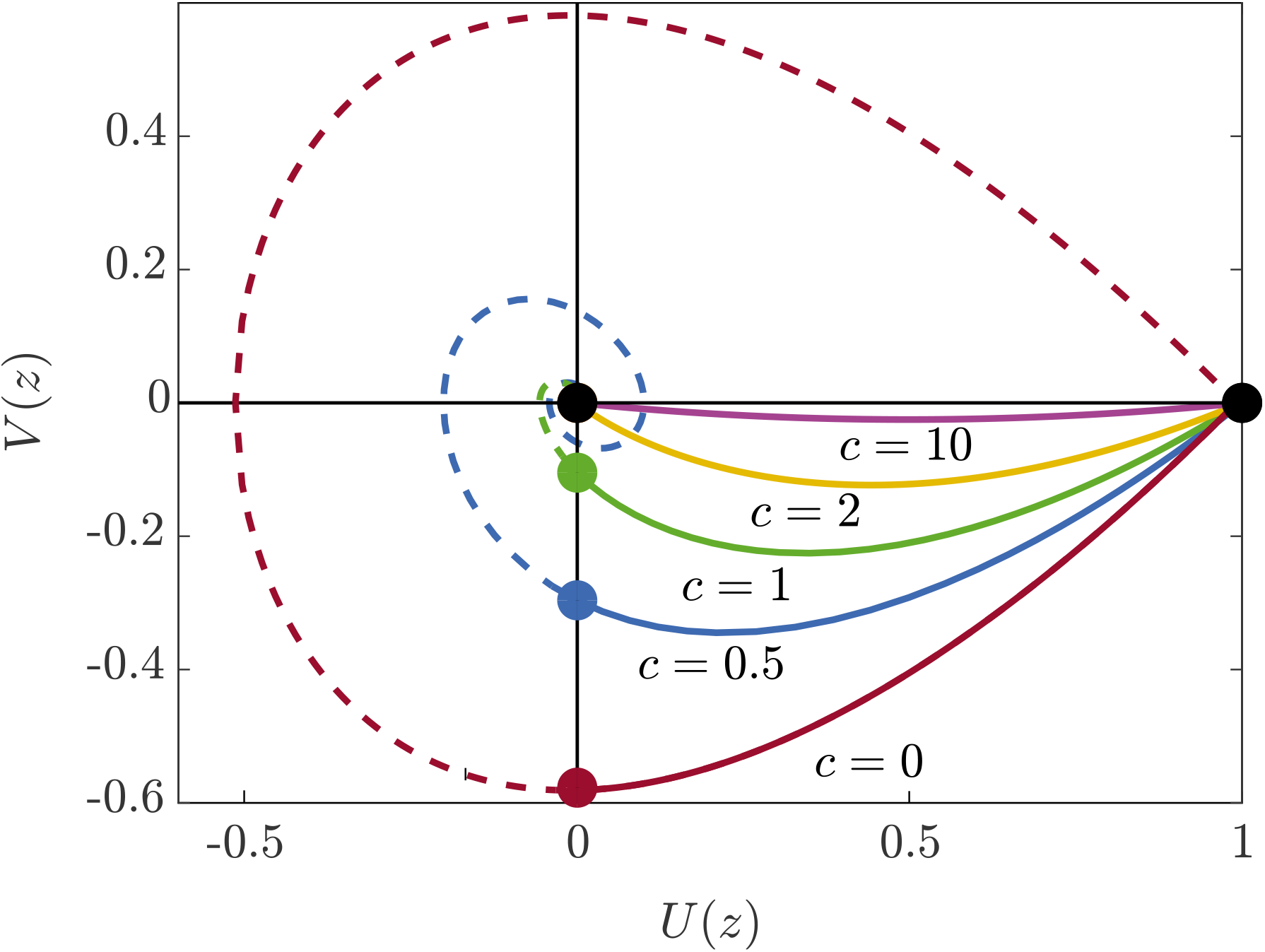
Family of trajectories in the phase plane for various choices of *c*. Heteroclinic trajectories running between (1, 0) to (0, 0) for *c* = 10, 2, 1 and 0.5, as indicated. An additional trajectory with *c* = 0 forms a homoclinic trajectory to (1, 0). Equilibrium points at (1, 0) and (0, 0) are shown with black discs. For the trajectories with 0 < *c* < 2 the intersection with the *U*(*z*) = 0 axis is shown with an appropriately coloured disc: green for *c* = 1; blue for *c* = 0.5; and red for *c* = 0.

**Figure 5:**
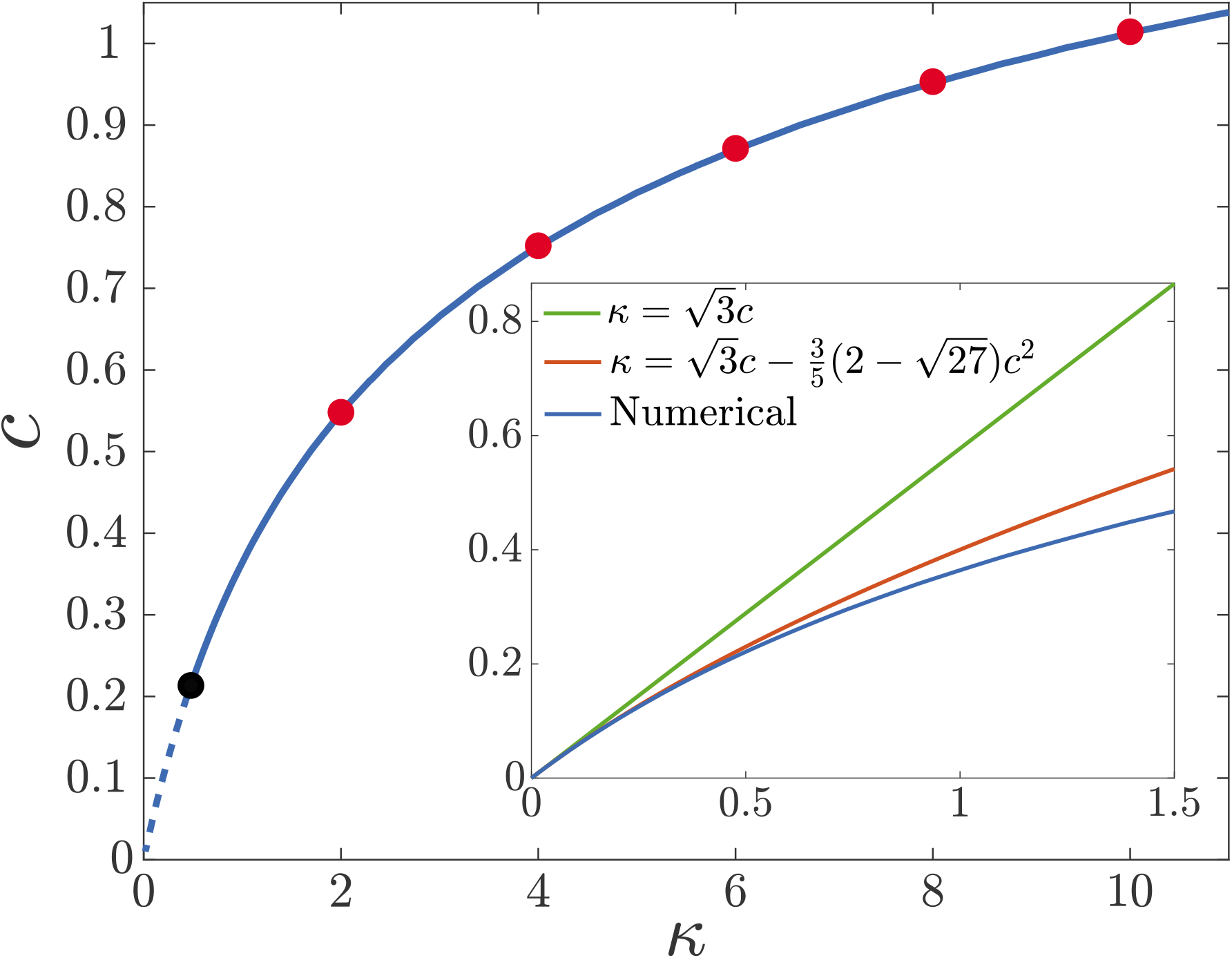
Relationship between *c* and *κ* for the Fisher-Stefan model. The blue curve is obtained by solving Equation (18) and calculating the value of *κ* corresponding to the intersection of (0, −*c/κ*). The red circles are obtained by solving Equations (2–4) and using the full time dependent solutions to estimate the eventual long time travelling wave speed, *c*. The black circle denotes the approximate critical value of *κ*_crit_. The Fisher-Stefan model is solved with Δ*ξ* = 1 × 10^−4^, Δ*t* = 1 × 10^−3^, *ϵ* = 1 × 10^−8^ and the initial condition given by Equation (8), where *α* = 0.5 and *β* = 1. The inset shows the comparison of the numerical solution the perturbation solutions. The green line represents the 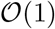 perturbation solution, and the red line represents the two-term 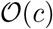 perturbation solution.

In addition to exploring the relationship between *κ* and *c* in the phase plane, we also solve the Fisher-Stefan model (9)-(12) numerically with a particular choice of initial condition given by Equation (8) with *β* = 1 and *α* = 0.5. We allow such numerical solutions to evolve for a sufficient duration of time that the resulting solution appears to settle into a travelling wave, from which we can estimate the speed, *c*. Repeating this procedure for various values of *κ* enables us to estimate how our choice of *κ* influences *c*. Additional results, shown as red discs in Figure 5, confirm that long time travelling wave solutions from the partial differential equation description compare very well with the relationship implied by the phase plane analysis.

As we stated in Section 2.2, whenever we are working in the phase plane we make the implicit assumption that a travelling wave solution has been generated. Yet, when we compare results in Figure 2(b)-(c) we know that long time travelling wave solutions do not always form, since this outcome depends upon the choice of *κ*. We explore this in Figure 5 by holding the initial condition constant in the numerical solution of Equations (9)–(12) and choosing a sequence of increasingly small values of *κ*. The numerical solutions suggest that for this initial condition there is a threshold value, *κ*_crit_ ≈ 0.48. If *κ* > *κ*_crit_ we observe long time travelling wave solutions and for *κ* < *κ*_crit_ the population eventually becomes extinct. This approximate threshold value is shown in Figure 5 as a black disc, and the relationship between *κ* and *c* obtained in the phase plane is shown as a solid line for *κ* > *κ*_crit_ and as a dashed line for *κ* < *κ*_crit_.

### 2.4 Critical length and the spreading-extinction dichotomy

We will now provide insight into the spreading-extinction dichotomy by examining time-dependent numerical solutions of Equations (9)–(12) in Figure 6. Figure 6(a) show numerical solutions for a particular choice of *κ* where we see some very interesting behaviour. At *t* = 30 the solution appears to be decreasing, almost to extinction, whereas by *t* = 60 and *t* = 90 the solution recovers from the initial decline to eventually form a travelling wave. Results in Figure 6(b) show details at intermediate values of *t* to clearly highlight this initial decline followed by the recovery. Figure 6(c) shows estimates of *L*(*t*) as a function of *t*, where we see that *L*(*t*) increases slowly at early time before eventually increasing at a constant rate, corresponding to a travelling wave solution. In contrast, Figure 6(d) shows the solution of Equations (9)–(12) for the same initial condition as in Figure 6(a) with the only difference being that *κ* is reduced. Figure 6(d) indicates that the population appears to be almost extinct at *t* = 90, and additional results magnified in Figure 6(e) show that the population never recovers, and instead goes extinct as t increases. Figure 6(f) shows *L*(*t*) as a function of *t*, where we see that the spreading population initially increases its domain before eventually stalling.

Du and colleagues [39] provide a formal proof of the existence of a critical length, *L*_crit_ = *π*/2, such that if ever *L*(*t*) > *L*_crit_ the population will evolve to a travelling wave, whereas if *L*(*t*) never exceeds this critical length then the population will eventually become extinct. Here we provide some simple physical and mathematical arguments to confirm this result. Visual inspection of the time dependent solutions of Equations (9)–(12) in Figure 6(b) and (e) confirm that we have *u*(*x,t*) ≪ 1 close to the time when population recovers (Figure 6(b)) or fails to recover from the initial decline (Figure 6(e)). This observation suggests that we can study the spreading-extinction dichotomy using an approximate linearised model where 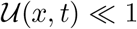. Under these conditions we can approximate the Fisher-Stefan model with

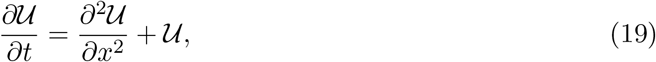

for 0 < *x* ≤ *L*, with 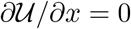 at *x* = 0 and 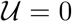 at *x* = *L*. In this approximate analysis we treat the domain length as a fixed quantity and this allows us to write down the exact solution of the linear equation (19) as

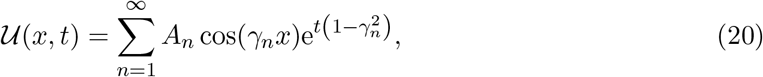

where *γ_n_* = *π*(2*n* − 1)/(2*L*), *n* = 1,2,3,…, and *A_n_* are constants chosen so that the solution matches the initial condition, 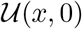. The solution of the linearised model (20) can be further simplified by assuming that the dynamics near the time of population recovery, or decline, can be approximated by taking a leading eigenvalue approximation so that

**Figure 6:**
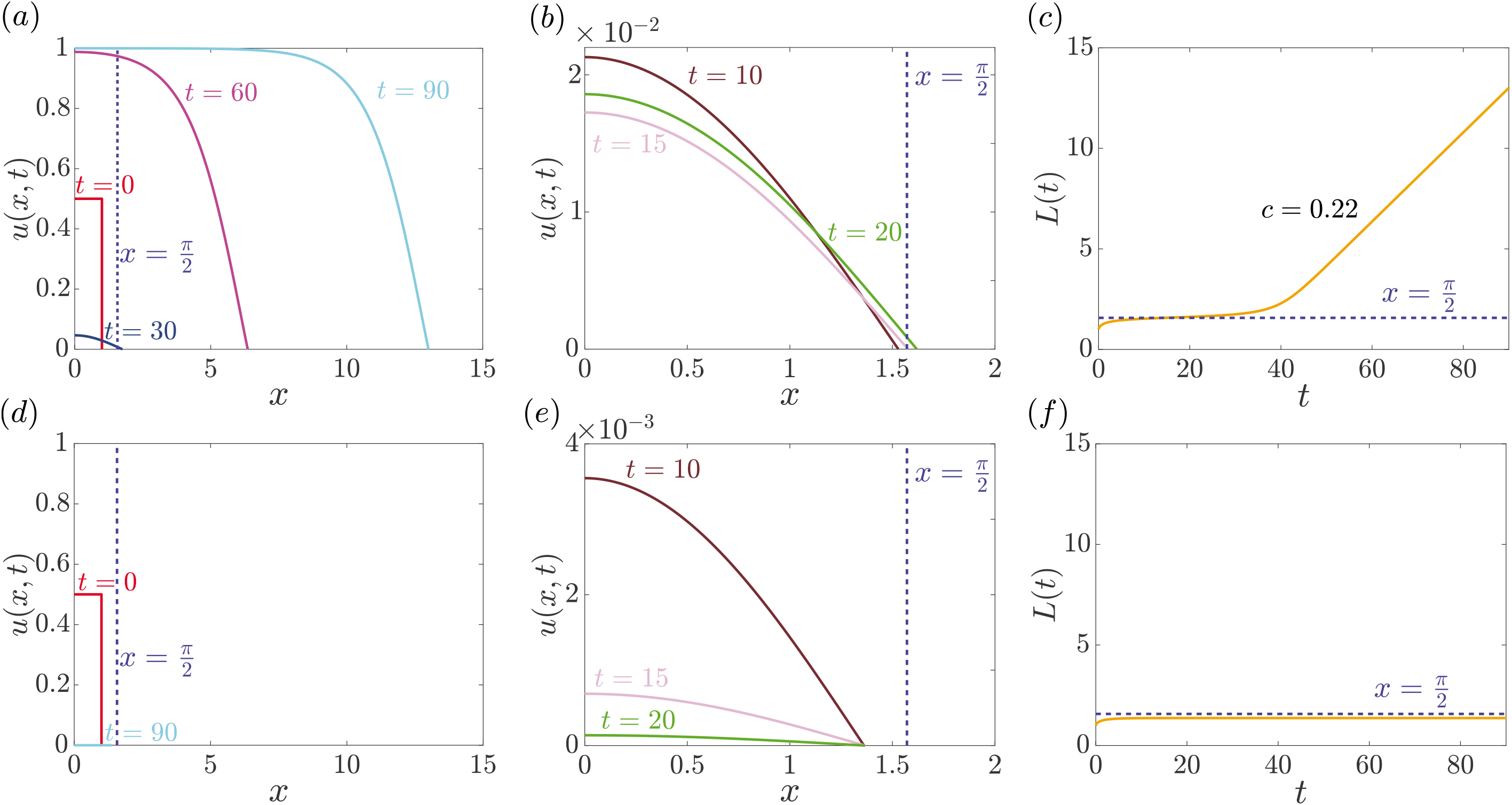
Numerical solutions of the Fisher-Stefan model showing both travelling wave and extinction phenomena. The first row represents the invasion phenomenon, and the second row represents the extinction. *(a)* Time evolution of the density profiles for invading cell population. *(b)* Magnified density profiles in *(a)* from *x* = 0 to 2 at intermediate times. *(c)* Progression of *L*(*t*) superimposed with the critical length of *π*/2. *(d)* Time evolution of the density profiles for extinction. *(e)* Magnified density profiles in *(d)* from *x* = 0 to 2 at intermediate times. *(f)* Progression of *L*(*t*) superimposed with the critical length of *π*/2. For both simulations, Δ*ξ* = 1 × 10^−4^, Δ*t* = 1 × 10^−3^, *ϵ* = 1 × 10^−8^, and the initial condition given by Equation (8), where *α* = 0.5 and *β* = 1. Results in *(a)-(c)* correspond to *κ* = 0.5, while results in *(d)-(f)* correspond to *κ* = 0.4.

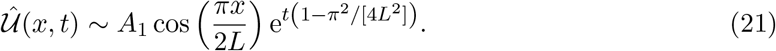

With this approximate solution we formulate a conservation statement describing the time rate of change of the total population within the domain,

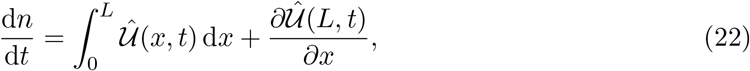

where 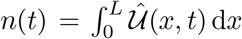 is the total population within the domain. The first term on the right of Equation (22) is the rate of increase of the total population owing to the source term and the second term on the right of Equation (22) is the rate of decrease of the total population owing to diffusive loss at the boundary at *x* = *L*. Setting d*n*/d*t* = 0, and substituting Equation (21) into (22) gives *L* = *L*_crit_ = *π*/2, corroborating the results of Du and colleagues [39]. We interpret this as follows. Once a time-dependent solution of Equations (9)–(12) evolves such that *L*(*t*) > *π*/2, the net positive accumulation of mass in the system means that a travelling wave solution will eventually form, as in Figure 6(a)-(c). Alternatively, if the time dependent solution of Equations (9)–(12) evolves such that *L*(*t*) never exceeds *π*/2, the net loss of mass in the system means that the population will always go extinct, as in Figure 6(d)-(f). It is also worthwhile to note that since the result that *L*_crit_ = *π*/2 is governed by a linearised model (19), this outcome will hold for any generalisation of the Fisher-Stefan model that can be linearised to give Equation (19). For example, if we extended the Fisher-Stefan model to consider a generalised logistic source term, *u*(1 − *u*)^*m*^, where *m* > 0 is some exponent [56, 57, 58, 59], then the same *L*_crit_ = *π*/2 would apply.

### 2.5 Perturbation solution when *c* ≪ 1

Now that we have used phase plane analysis and linearisation to establish conditions for travelling wave solutions of the Fisher-Stefan model to form, we turn our attention to whether it is possible to provide additional mathematical insight into the details of the shape of the travelling wave solutions. It is well known that travelling wave solutions of the Fisher-KPP model travel with speed *c* ≥ *c*_min_ = 2, and that it is possible to obtain approximate perturbation solutions to describe the shape of the travelling wave solutions in large *c* limit [4, 5, 31]. We now attempt to follow a similar analysis to describe the shape of the travelling wave solution of the Fisher-Stefan model for which *c* < 2, suggesting that we attempt to find a perturbation solution for small *c*:

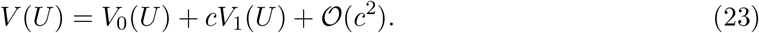

Substituting this expansion into Equation (18) we obtain ordinary differential equations governing *V*_0_(*U*) and *V*_1_(*U*) which can be integrated exactly. Ensuring that *U*(1) = 0 we obtain

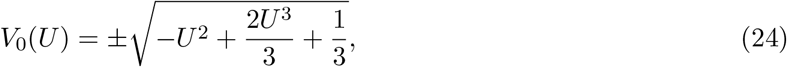

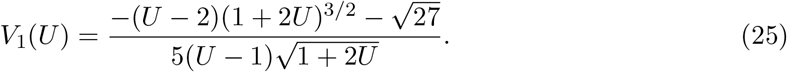

Since we solve for both *V*_0_(*U*) and *V*_1_(*U*) we can construct both an 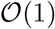 perturbation solution given by 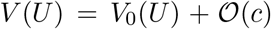, as well as an 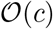 perturbation solution given by 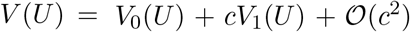. To compare the accuracy of these perturbation solutions for the shape of the *V*(*U*) curve in the phase plane we generate a series of numerical phase planes for *c* = 0.05,0.1, 0.2 and 0.5 in Figure 7. The numerical trajectories, shown in blue, run between the equilibrium points (1, 0) and (0, 0), and pass through the point (0, −*c/κ*). In the numerical solutions we highlight (0, −*c/κ*) with a blue disc. In each subfigure of Figure 7, we compare the numerical trajectory with the 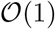 perturbation solution in red. In each case the 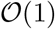 perturbation solution runs between (1, 0) and first intersects the *V*-axis at 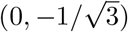. We show this point with a red disc. Comparing the red and blue trajectories in the fourth quadrant clearly shows a discrepancy that increases with *c*, as expected. Similarly, in each subfigure of Figure 7 we also compare the numerical trajectory with the 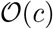 perturbation solution shown in green. In each case the 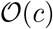 perturbation solution runs between (1, 0) and first intersects the *V*-axis at 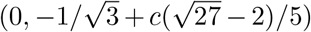, and we show this point with a green disc. Comparing the green and blue trajectories in the fourth quadrant shows that we have an excellent match between the numerical and perturbation solutions. This comparison indicates that the 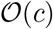 perturbation solution can be used to provide a highly-accurate approximation of the shape of the travelling wave solutions of the Fisher-Stefan model for *c* < 0.5.

Using Equations (23)–(25) we can obtain additional analytical insight into the relationship between *c* and *κ* that we previously explored numerically in Figure 5. Since the ordinate of the intersection point is *V* = −*κ/c*, we can develop approximate closed-form relationships between *c* and *κ*. These relationships are plotted in the inset of Figure 5 for *κ* < 2 and *c* < 1. Comparing the numerically deduced relationship between *c* and *κ* with the perturbation solutions shows a good match, with the expected result that the 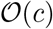 perturbation solution leads to a particularly accurate approximation.

**Figure 7:**
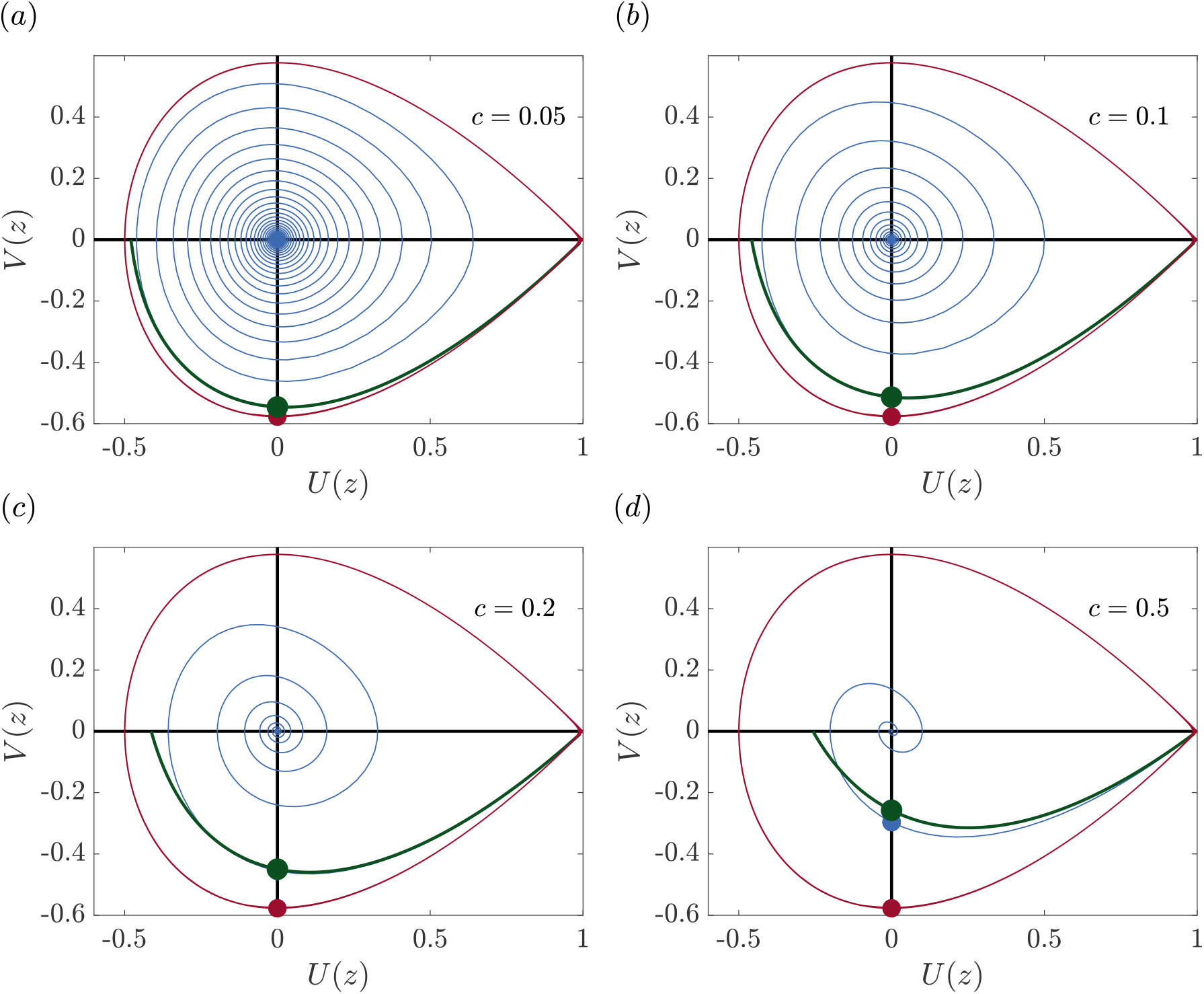
Comparison of numerical trajectories in the phase plane and perturbation solutions for various choices of *c*. The blue lines are the numerical trajectories for *c* = 0.05,0.1,0.2 and 0.5 in *(a)-(d)*, respectively. The red curves are the 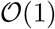 perturbation solution. The green curves are the 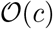 perturbation solution. The red, blue and green discs indicate the intersection points of the trajectories with *U*(*z*) = 0.

Since our perturbation solutions provide good approximations to the *V*(*U*) curve in the fourth quadrant of the phase plane, shown in Figure 7, we now compare the accuracy of the perturbation solutions in the travelling wave coordinate system. For the 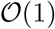 perturbation solution we have

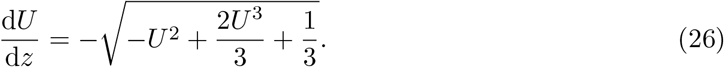

Integrating Equation (26) with *U* = 0 at *z* = 0 gives an implicit solution

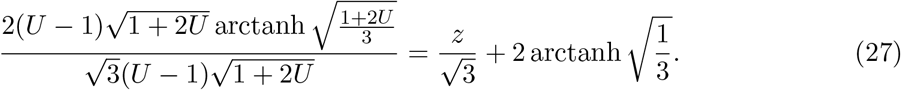

For the 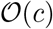 perturbation solution we have

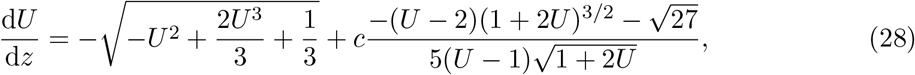

for which we cannot find an exact solution. Nonetheless, Equation (28) can be integrated numerically to give a numerical approximation of *U*(*z*).

Figure 8 shows a suite of travelling wave solutions obtained by solving Equations (9)–(12) (in dashed blue) presented for *c* = 0.05, 0.1,0.2 and 0.5. In each case the solutions are obtained for a sufficiently long time that the full time dependent numerical solutions have settled into a travelling wave. These travelling waves are then shifted so that *u*(*x, t*) = 0 at *z* = 0, where *z* = *x−ct*. For each value of *c*, we superimpose plots of the 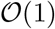 solution, given by Equation (27) (in solid red). The results show that the 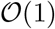 solution provides an excellent match to the shape of the full numerical solutions of Equations (9)–(12) for *c* = 0.05. For larger *c*, however, the 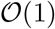 solution provides a relatively poor approximation. Similarly, for each value of *c* we also plot the 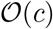 solution (in solid green). Here we see that the 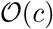 perturbation solution provides an excellent match, being indistinguishable from the full numerical solutions of Equations (9)–(12) for *c* < 0.1. In each subfigure of Figure 8 we also provide an inset showing a magnified region just behind the leading edge of the wavefront to make the comparison between the numerical solution of the partial differential equation and the two perturbation solutions clearer.

## 3 Conclusion

In this work we directly compare features of the travelling wave solutions of the well-known Fisher-KPP model and solutions to the Fisher-Stefan model. A key feature of the Fisher-KPP model is that any positive initial condition with compact support will always evolve into a travelling wave that moves with the minimum wave speed, *c*_min_ = 2. Therefore, according to the Fisher-KPP model, any initial population will lead to successful invasion. This feature is a potential weaknesses of the Fisher-KPP model since it is well known that small translocated populations do not always invade, and can become extinct [46]. In contrast, the Fisher-Stefan model is an adaptation of the Fisher-KPP model with a moving boundary, *x* = *L*(*t*). In the Fisher-Stefan model, the evolution of the moving boundary is governed by a one-phase Stefan condition [39, 40, 41, 42, 43, 44, 45]. The Fisher-Stefan model can support travelling wave solutions with speed *c* < 2. Since both the Fisher-KPP and the Fisher-Stefan model support travelling wave solutions, both of these models can be used to simulate invasion processes. However, unlike the Fisher-KPP model, the Fisher-Stefan model also predicts the extinction of certain initial populations, giving rise to the spreading-extinction dichotomy [39, 40, 41, 42, 43, 44, 45].

The spreading-extinction dichotomy associated with the Fisher-Stefan model has been studied, mainly using rigorous mathematical approaches, leading to many important existence results [39, 40, 41, 42, 43, 44, 45]. One of the aims of this work is to provide a more practical comparison of the Fisher-KPP and Fisher-Stefan models using standard tools of applied mathematics and engineering to provide insight into the similarities and differences between these two models of invasion. It is interesting to note that the partial differential equation descriptions of the Fisher-KPP and Fisher-Stefan models are very different, since the Fisher-KPP model is associated with a fixed domain and the Fisher-Stefan model is a moving boundary problem. In contrast, when we analyse the travelling wave solutions of both models we find that the equations governing the trajectories in the phase plane are the same. Interestingly, standard phase plane arguments for the Fisher-KPP model lead us to conclude that travelling wave solutions with speed less than the minimum *c*_min_ = 2 are not possible based on physical arguments and therefore not normally recorded. In this work we show that the travelling wave solutions of the Fisher-Stefan model require that *c* < 2, and it turns out that it is precisely these normally-discarded solutions for the Fisher-KPP model that are relevant for the Fisher-Stefan model.

**Figure 8:**
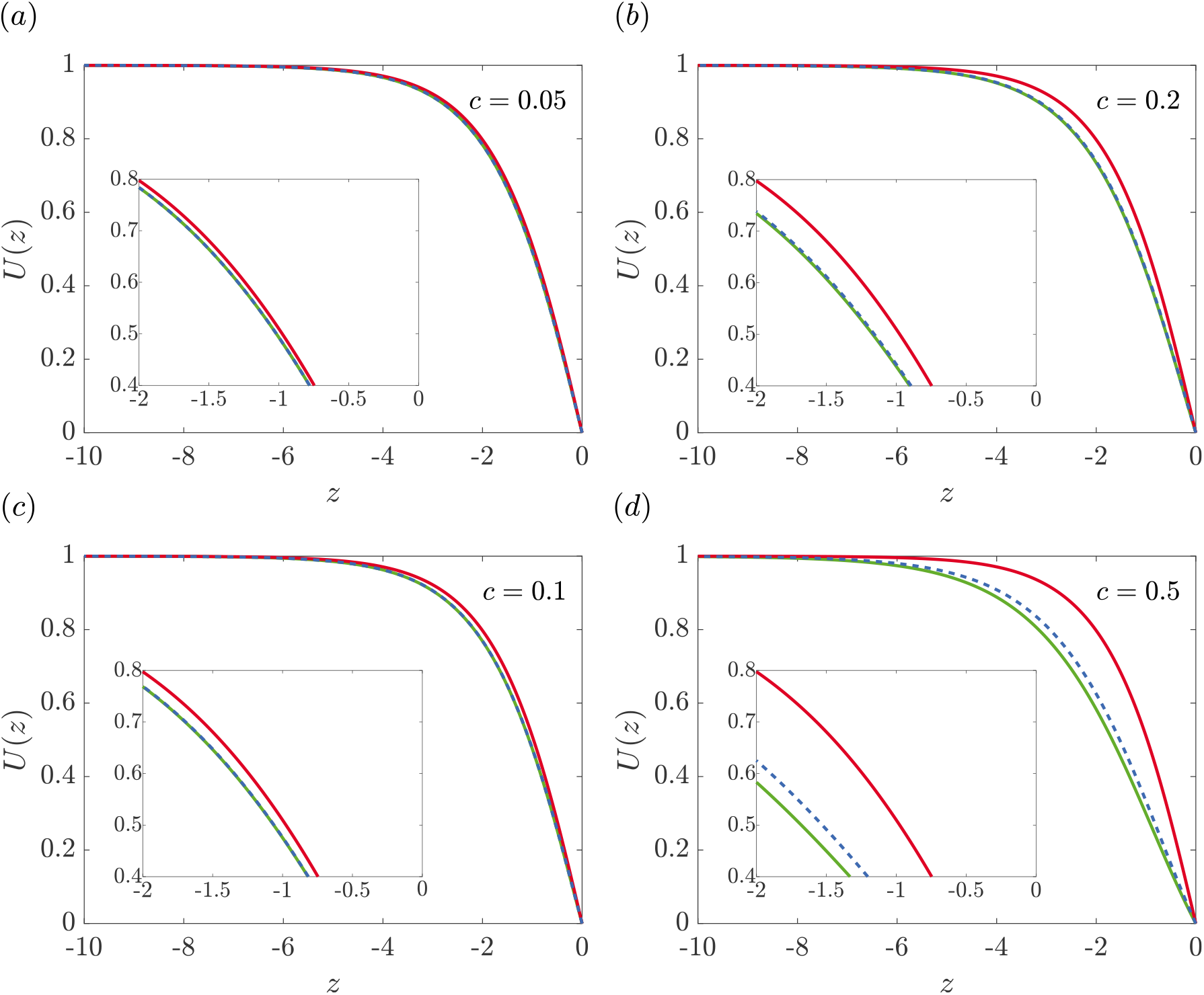
Density profiles comparing the numerical solution of the Fisher-Stefan equation with the perturbation solution for *c* ≪ 1. *(a)* c = 0.05. *(b)* c = 0.1. *(c)* c = 0.2. *(d)* c = 0.5. The solid blue line represents the travelling wave solution obtained from the time-dependent Fisher-Stefan model, (2) and shifting the resulting travelling wave profile so that *U*(0) = 0. The red solid line represents the travelling wave profile obtained from the 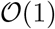 perturbation solution and the dashed green line represents the travelling wave profile obtained from the 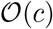 perturbation solution. In each subfigure we show an inset magnifying the travelling wave profiles so that the differences between the numerical and perturbation solutions are visually distinct.

In the non-dimensional Fisher-Stefan model there is one free parameter, *κ* > 0, that relates the dynamics of the moving boundary, *x* = *L*(*t*), and the spatial gradient of the density function at the moving boundary. By analysing trajectories in the phase plane associated with travelling wave solutions of the Fisher-Stefan model we are able to arrive at a relationship between *κ* and *c*, confirming that *c* → 2^−^ as *κ* → ∞. However, all phase plane analysis of the Fisher-Stefan model implicitly assumes that a travelling wave solution has formed, whereas numerical solutions of the full partial differential equation description of the Fisher-Stefan model shows that for a fixed initial condition there is a critical value *κ*_crit_: for *κ* > *κ*_crit_ the solution eventually evolves to a travelling wave, whereas for *κ* < *κ*_crit_ the solution eventually goes extinct. The time-dependent solutions of the partial differential equation models suggest that near this transition between eventual extinction and eventual travelling wave formation, we have *u*(*x,t*) ≪ 1, suggesting that we can obtain insight using a linearised model. Working in a linearised framework we obtain an approximate solution from which we form a conservation statement describing the net accumulation of total population numbers in the domain. In the critical case where there is zero net accumulation of mass in the domain, we find that there is a critical length, *L*_crit_ = *π*/2, and whenever *L*(*t*) exceeds *π*/2 the solution will always evolve to a travelling wave while if *L*(*t*) never exceeds *π*/2 the density will always eventually go extinct. Using a comparison with the Stefan problem, Equations (9)–(12) with 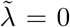, we can strengthen these results to be that if *L*(*t*) exceeds

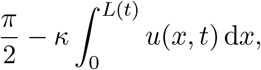

then a travelling wave will form. Or, if the population goes extinct with *L*(*t*) → *L*_e_ as *t* → ∞, then

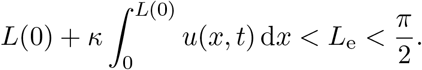

While it is well-known that there are no closed-form exact solutions describing travelling wave solutions of the Fisher-KPP equation for arbitrary *c*, it is possible to obtain approximate perturbation solutions for *c* ≫ 1 [4, 5]. Since travelling wave solutions for the Fisher-Stefan model move with speed *c* < 2, we construct a perturbation solution for *c* ≪ 1, leading to approximate closed form solutions for the shape of the trajectory in the phase plane which can be used to estimate the shape of the travelling wave. We find that the 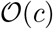 perturbation solutions provide an excellent match to our full numerical solutions for *c* < 0.5, thereby providing analytical insight into the relationship between the speed and shape of the travelling wave solutions of the Fisher-Stefan model.

The purpose of this work is to compare the Fisher-Stefan and Fisher-KPP models of invasion. Although we begin our work by pointing out that Fisher-Stefan model enables us to simulate population extinction, whereas Fisher-KPP does not, there are also limitations of the Fisher-Stefan model that warrant acknowledgement and discussion. For example, time-dependent solutions of the Fisher-Stefan model that move in the positive *x*-direction, including travelling wave solutions, involve a loss of the population at the free boundary, *x* = *L*(*t*), since *∂u/∂x* < 0 at *x* = *L*(*t*). It could be difficult to justify this loss at the moving boundary based on biological, ecological or physical grounds and/or to calibrate the model to estimate a relevant value of *κ*. To address this point, it is worthwhile recalling that the Fisher-Stefan model makes use of a one-phase Stefan boundary condition which is a simplification of a more realistic two-phase Stefan boundary condition [47, 48]. In more realistic applications of invasion, such as malignant cellular invasion into surrounding tissues, there will be a conversion of consumed tissue into malignant cells at the interface [34, 60, 61, 62]. One way of interpreting this conversion from tissues to cells is that there is a loss of one species, in this case the surrounding tissue, that is converted into another species, in this case invasive cells. Therefore, while we appreciate that the practical interpretation of loss at the moving boundary in the one phase Fisher-Stefan model is difficult to justify, we anticipate that this loss at a moving boundary would be very natural in an extended framework where the Fisher-Stefan model is re-cast as a two-phase problem. `Data Access` This article has no additional data. Key algorithms used to generate results are available on Github at `GitHub`.

## References

[1] Aronson DG, Weinberger HF. 1978. Multidimensional nonlinear diffusion arising in population genetics. Adv. Math. 30, 33–76. (doi:10.1016/0001-8708(78)90130-5).

[2] Fisher RA. 1937. The wave of advance of advantageous genes. Ann. Eugenic. 7, 355–369. (doi:10.1111/j.1469-1809.1937.tb02153.x).

[3] Kolmogorov AN, Petrovskii IG, Piskunov NS, 1937. A study of the diffusion equation with increase in the amount of substance, and its application to a biological problem. Bull. Moscow Univ. Math. Mech. 1, 1–26.

[4] Canosa J. 1973. On a nonlinear diffusion equation describing population growth. IBM J. Res. Dev. 17, 307–313. (doi:10.1147/rd.174.0307).

[5] Murray JD. 2002. Mathematical biology I. An introduction New York: Springer.

[6] Grindrod P. 2007. Patterns and waves Oxford: Oxford University Press.

[7] Maini PK, McElwain DLS, Leavesley DI. 2004. Traveling wave model to interpret a wound-healing cell migration assay for human peritoneal mesothelial cells. Tissue Eng. 10, 475–482. (doi:10.1089/107632704323061834).

[8] Maini PK, McElwain DLS, Leavesley D. 2004. Travelling waves in a wound healing assay. Appl. Math. Lett. 17, 575–580. (doi:10.1016/S0893-9659(04)90128-0).

[9] Simpson MJ, Treloar KK, Binder BJ, Haridas P, Manton KJ, Leavesley DI, McElwain DLS, Baker RE. 2013. Quantifying the roles of cell motility and cell proliferation in a circular barrier assay. J. R. Soc. Interface. 10, 20130007. (doi:10.1098/rsif.2013.0007).

[10] Sherratt JA, Murray JD. 1990. Models of epidermal wound healing. Proc. R. Soc. Lond. B. 241, 29–36. (doi:10.1098/rspb.1990.0061).

[11] Skellam JG. 1951. Random dispersal in theoretical populations. Biometrika 38, 196–218. (doi:10.1093/biomet/38.1-2.196).

[12] Shigesada N, Kawasaki K, Takeda Y. 1995. Modeling stratified diffusion in biological invasions. Am. Nat. 146, 229–251. (doi:10.1086/285796).

[13] Steele J, Adams J, Sluckin T. 1998. Modelling paleoindian dispersals. World Archaeol. 30, 286–305. (doi:10.1080/00438243.1998.9980411).

[14] Levin SA, Muller-Landau HC, Nathan R, Chave J. 2003. The ecology and evolution of seed dispersal: a theoretical perspective. Annu. Rev. Ecol. Syst. 34, 575–604. (doi:10.1146/annurev.ecolsys.34.011802.132428).

[15] Cai AQ, Landman KA, Hughes BD. 2007. Multi-scale modeling of a wound-healing cell migration assay. J. Theor. Biol. 245, 576–594. (doi:/10.1016/j.jtbi.2006.10.024).

[16] Sengers BG, Please CP, Oreffo ROC. 2007. Experimental characterization and computational modelling of two-dimensional cell spreading for skeletal regeneration. J. R. Soc. Interface 4, 1107–1117. (doi:10.1098/rsif.2007.0233).

[17] Tremel A, Cai A, Tirtaatmadja N, Hughes BD, Stevens GW, Landman KA, O’Connor AJ. 2009. Cell migration and proliferation during monolayer formation and wound healing. Chem. Eng. Sci. 64, 247–253. (doi:10.1016/j.ces.2008.10.008).

[18] Nardini JT, Chapnick DA, Liu X, Bortz DM. 2016. Modeling keratinocyte wound healing dynamics: Cell–cell adhesion promotes sustained collective migration. J. Theor. Biol. 400, 103–117. (doi:10.1016/j.jtbi.2016.04.015).

[19] Warne DJ, Baker RE, Simpson MJ. 2019. Using experimental data and information criteria to guide model selection for reaction–diffusion problems in mathematical biology. B. Math. Biol. 81, 1760–1804. (doi:10.1007/s11538-019-00589-x).

[20] Swanson KR, Bridge C, Murray JD, Alvord EC Jr. 2003. Virtual and real brain tumors: using mathematical modeling to quantify glioma growth and invasion. J. Neurol. Sci. 216, 1–10. (doi:/10.1016/j.jns.2003.06.001).

[21] Swanson KR, Rostomily RC, Alvord EC Jr. 2008. A mathematical modelling tool for predicting survival of individual patients following resection of glioblastoma: a proof of principle. Brit. J. Cancer. 98, 113–119. (doi:/10.1038/sj.bjc.6604125).

[22] Pérez-Beteta J, Martínez-González A, Pérez-García VM. 2018. A three-dimensional computational analysis of magnetic resonance images characterizes the biological aggressiveness in malignant brain tumours. J. R. Soc. Interface 15, 20180503. (doi:10.1098/rsif.2018.0503).

[23] Mercer GN, Weber RO. 1995. Combustion wave speed. Proc. R. Soc. A-Math. Phy. 450, 193–198. (doi:10.1098/rspa.1995.0079).

[24] Tang S, Qin S, Weber R. 1993. Numerical studies on 2–dimensional reaction–diffusion equations. J. Aust. Math. Soc. B. 35, 223–243. (doi:10.1017/S0334270000009140).

[25] Forbes L. 1997. A two-dimensional model for large–scale bushfire spread. J. Aust. Math. Soc. B. 39, 171–194. (doi:10.1017/S0334270000008791).

[26] Treloar KK, Simpson MJ, McElwain DLS, Baker RE. 2014. Are *in vitro* estimates of cell diffusivity and cell proliferation rate sensitive to assay geometry? J. Theor. Biol. 356, 71–84. (doi:10.1016/j.jtbi.2014.04.026).

[27] Jin W, Lo K-Y, Chou S-E, McCue SW, Simpson MJ. 2018. The role of initial geometry in experimental models of wound closing. Chem. Eng. Sci. 179, 221–226. (doi:10.1016/j.ces.2018.01.004).

[28] King JR, McCabe PM. 2003. On the Fisher–KPP equation with fast nonlinear diffusion. Proc. R. Soc. A-Math. Phy. 459, 2529–2546. (doi:/10.1098/rspa.2003.1134).

[29] Sánchez-Garduño F, Maini PK 1994. Existence and uniqueness of a sharp travelling wave in degenerate non-linear diffusion Fisher-KPP equations, J. Math. Bio. 33, 163–192. (doi:10.1007/BF00160178).

[30] McCue SW, Jin W, Moroney TJ, Lo K-Y, Chou S-E, Simpson MJ. 2019. Hole-closing model reveals exponents for nonlinear degenerate diffusivity functions in cell biology. To appear, Physica. D. arXiv:1903.10800 [physics.bio-ph].

[31] Simpson MJ, Landman KA, Hughes BD, Newgreen DF. 2006. Looking inside an invasion wave of cells using continuum models: Proliferation is the key. J. Theor. Biol. 243, 343–360. (doi:10.1016/j.jtbi.2006.06.021).

[32] Witelski TP. 1995. Merging traveling waves for the porous-Fisher’s equation. Appl. Math. Lett. 8, 57–62. (doi:10.1016/0893-9659(95)00047-T).

[33] Sherratt, JA, Marchant BP. 1996. Non-sharp travelling wave fronts in the Fisher equation with degenerate nonlinear diffusion. Appl. Math. Lett. 9, 33–38. (doi:10.1016/0893-9659(96)00069-9).

[34] Perumpanani AJ, Sherratt JA, Norbury J, Byrne H. 1999. A two parameter family of travelling waves with a singular barrier arising from the modelling of matrix mediated malignant invasion. Physica D. 126, 145–159. (doi:10.1016/S0167-2789(98)00272-3).

[35] Marchant BP, Norbury J, Sherratt JA. 2001. Travelling wave solutions to a haptotaxis-dominated model of malignant invasion. Nonlinearity 14, 1653–1671. (doi:10.1088/0951-7715/14/6/313).

[36] Landman KA, Simpson MJ, Slater JA, Newgreen DF. 2005. Diffusive and chemotactic cellular migration: smooth and discontinuous travelling wave solutions. SIAM J. Appl. Math. 65, 1420–1442. (doi:10.1137/040604066).

[37] Painter KJ, Sherratt JA. 2003. Modelling the movement of interacting cell populations. J. Theor. Biol. 225, 327–339. (doi:10.1016/j.jtbi.2015.01.025).

[38] Painter KJ, Hillen T. 2013. Mathematical modelling of glioma growth: the use of diffusion tensor imaging (DTI) data to predict the anisotropic pathways of cancer invasion. J. Theor. Biol. 323, 25–39. (doi:10.1016/j.jtbi.2013.01.014).

[39] Du Y, Lin Z. 2010. Spreading–vanishing dichotomy in the diffusive logistic model with a free boundary. SIAM J. Math. Anal. 42, 377–405. (doi:10.1137/090771089).

[40] Du Y, Guo Z. 2011. Spreading–vanishing dichotomy in a diffusive logistic model with a free boundary, II. J. Differ. Equations. 250, 4336–4366. (doi:10.1016/j.jde.2011.02.011).

[41] G Bunting, Y Du, K Krakowski. 2012. Spreading speed revisited: Analysis of a free boundary model. Netw. Heterog. Media 7, 583–603. (doi:10.3934/nhm.2012.7.583).

[42] Du Y, Guo Z. 2012. The Stefan problem for the Fisher-KPP equation. J. Differ. Equations. 253, 996–1035. (doi:10.1016/j.jtbi.2006.06.021).

[43] Du Y, Matano H, Wang K. 2014. Regularity and asymptotic behavior of nonlinear Stefan problems. Arch. Rational Mech. Anal. 212, 957–1010. (doi:10.1007/s00205-013-0710-0).

[44] Du Y, Matsuzawa H, Zhou M. 2014. Sharp estimate of the spreading speed determined by nonlinear free boundary problems. SIAM J. Math. Anal. 46, 375–396. (doi:10.1137/130908063).

[45] Du Y, Lou B. 2015. Spreading and vanishing in nonlinear diffusion problems with free boundaries. J. Eur. Math. Soc. 17, 2673–2724. (doi:10.4171/JEMS/568).

[46] Griffith B, Michael Scott J, Carpenter JW, Reed C. 1989. Translocation as a species conservation tool: status and strategy. Science 245, 477–480. (doi: 10.1126/science.245.4917.477).

[47] Crank J. 1987. Free and moving boundary problems. Oxford University Press, Oxford.

[48] Gupta SC. 2017. The classical Stefan problem. Basic concepts, modelling and analysis with quasi-analytical solutions and methods. Second edition. Elsevier, Amsterdam.

[49] Crank J. 1979. The mathematics of diffusion. Oxford University Press, Oxford.

[50] King JR, Riley DS, Wallman AM. 1999. Two-dimensional solidification in a corner. Proc. R. Soc. A-Math. Phy. 455, 3449–3470. (doi:10.1098/rspa.1999.0460).

[51] McCue SW, King JR, Riley DS. 2003. Extinction behaviour for two-dimensional inward-solidification problems. Proc. R. Soc. A-Math. Phy. 459, 977–999. (doi:10.1098/rspa.2002.1059).

[52] McCue SW, King JR, Riley DS. 2005. The extinction problem for three-dimensional inward solidification. J. Eng. Math. 52, 389–409. (doi:10.1007/s10665-005-3501-2).

[53] McCue SW, Wu B, Hill JM. 2008. Classical two-phase Stefan problem for spheres. Proc. R. Soc. A-Math. Phy. 464, 2055–2076. (doi:10.1098/rspa.2007.0315).

[54] King JR, Riley DS. 2000. Asymptotic solutions to the Stefan problem with a constant heat source at the moving boundary. Proc. R. Soc. A-Math. Phy. 456, 1163–1174. (doi:10.1098/rspa.2000.0556).

[55] Harley K, van Heijster P, Marangell R, Pettet GJ, Wechselberger M. 2015. Numerical computation of an Evans function for travelling waves. Math. Biosci. 266, 36–51. (doi:10.1016/j.mbs.2015.05.009).

[56] Simpson MJ, Landman KA, Hughes BD. 2010. Cell invasion with proliferation mechanisms motivated by time-lapse data. Physica A. 389, 3779–3790. (doi:10.1016/j.physa.2010.05.020).

[57] Tsoularis A, Wallace J. 2002. Analysis of logistic growth models. Math. Biosci. 179, 21–55. (doi:10.1016/S0025-5564(02)00096-2).

[58] Broadbridge P, Bradshaw BH, Fulford GR, Aldis GK. 2002. Huxley and Fisher equations for gene propagation: An exact solution. ANZIAM J. 44, 11–20. (doi:10.1017/S1446181100007860).

[59] Bradshaw-Hajek BH, Broadbridge P. 2004. A robust cubic reaction-diffusion system for gene propagation. Math. Comput. Model. 39, 1151–1163. (doi:10.1016/S0895-7177(04)90537-7).

[60] Gatenby RA, Gawlinski ET. 1996. A reaction-diffusion model of cancer invasion. Cancer Res. 56, 5745–5753. (doi:10.1007/s00285-013-0665-7).

[61] Haridas P, McGovern JA, McElwain DLS, Simpson MJ. 2017. Quantitative comparison of the spreading and invasion of radial growth phase and metastatic melanoma cells in a three-dimensional human skin equivalent model. PeerJ. 5, e3754. (doi:10.7717/peerj.3754).

[62] Browning AP, Haridas P, Simpson MJ. 2019. A Bayesian sequential learning framework to parameterise continuum models of melanoma invasion into human skin. B. Math. Biol. 81, 676–698. (doi:10.1007/s11538-018-0532-1).

