## Supplementary Information for "Revisiting the Fisher-KPP equation to interpret the spreading-extinction dichotomy"

June 17, 2019

[1] School of Mathematical Sciences, Queensland University of Technology (QUT), Brisbane, Australia.

[2] School of Science and Technology, University of New England, Armidale, Australia.

**Keywords**: Reaction-diffusion; Travelling wave; Invasion; Extinction; Moving boundary problem; Stefan problem.

### 1 Numerical methods

#### 1.1 Fisher-KPP model

We solve the Fisher-KPP equation

$$\frac{\partial u}{\partial t} = \frac{\partial^2 u}{\partial x^2} + u(1-u), \tag{1}$$

on $0 \leq x \leq x_\infty$, with $x_\infty$ chosen to be sufficiently large. We discretise the domain with a uniform finite difference mesh, with spacing $\Delta x$. We approximate the spatial derivatives in Equation (1) using a central finite difference approximation, and we integrate Equation (1) using an implicit Euler approximation, giving rise to

$$\frac{u_i^{j+1} - u_i^j}{\Delta t} = \left( \frac{u_{i-1}^{j+1} - 2u_i^{j+1} + u_{i+1}^{j+1}}{\Delta x^2} \right) + u_i^{j+1}(1 - u_i^{j+1}), \tag{2}$$

for $i = 2, \ldots, m-1$, where $m = x_\infty/\Delta x + 1$ is the total number of spatial nodes on the finite difference mesh, and the index $j$ represents the time index so that we approximate $u(x,t)$ by $u_i^j$, where $x = (i-1)\Delta x$ and $t = j\Delta t$.

For all numerical solutions of Equation (1) we enforce no-flux boundary conditions at $x = 0$ and $x = x_\infty$

$$u_2^{j+1} - u_1^{j+1} = 0, \tag{3}$$

$$u_m^{j+1} - u_{m-1}^{j+1} = 0. \tag{4}$$

Together, Equations (2)–(4) form a nonlinear system of algebraic equations that describe how to approximate $u_i^{j+1}$ from $u_i^j$ for $i = 1, \ldots, m$.

We use Newton-Raphson iteration to solve this non-linear system and we continue with the iterations until the infinity norm of the difference between successive estimates of $u_i^{j+1}$ falls below some small tolerance, $\epsilon$. For all results presented we are always careful to choose $\Delta x$, $\Delta t$ and $\epsilon$ so that the numerical algorithm produces grid-independent results. To illustrate the accuracy of our algorithm we present results in Figure 1 showing the evolution of the solution of Equation (1) evolving from an initial condition with compact support. Here we see that the solution rapidly approaches a constant shape, constant speed travelling wave what moves with the minimum wave speed, $c = 2$, as expected [1].

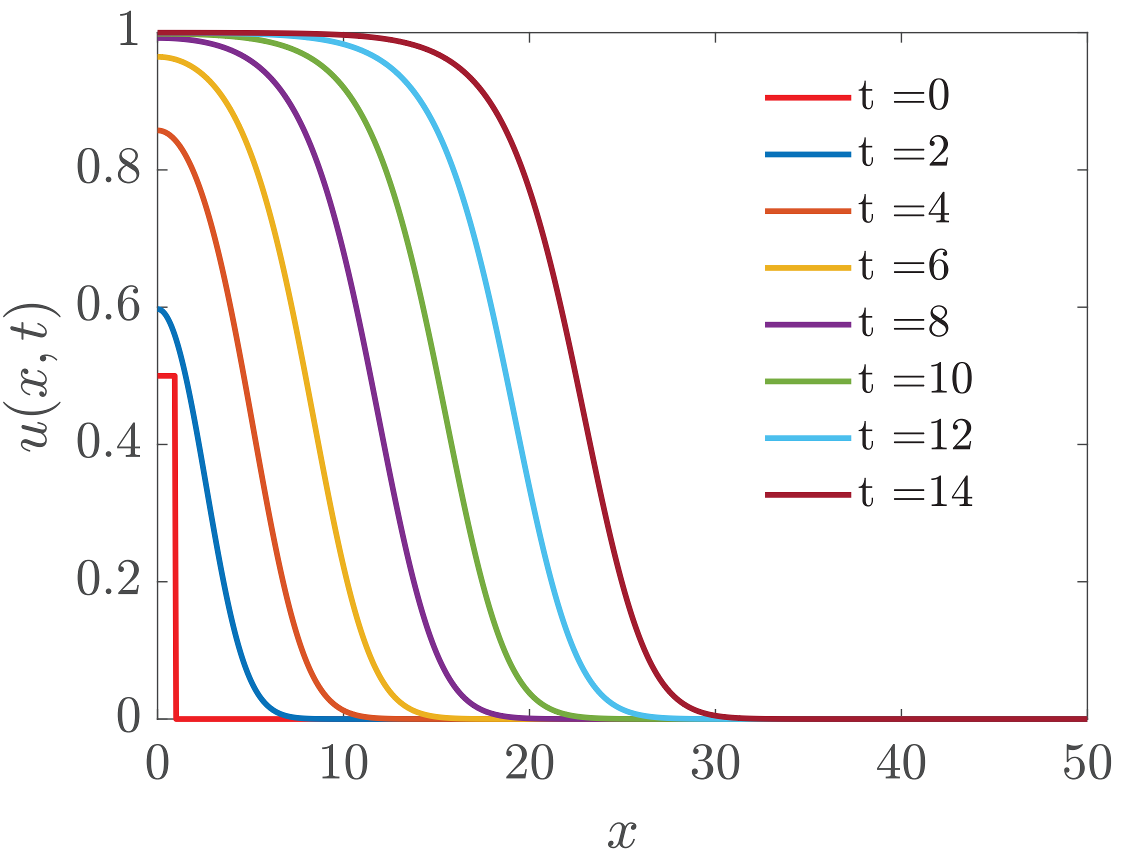


Figure 1: Numerical solutions of Equation (1) with $\Delta x = 1 \times 10^{-4}$, $\Delta t = 1 \times 10^{-3}$, $x_\infty = 50$ and $\epsilon = 1 \times 10^{-8}$. The initial condition is $u(x, 0) = 0.5$ for $x < 1$ and $u(x, 0) = 0.5$ for $x > 1$.

We have confidence in our numerical results in Figure 1 since we find that the results are grid-independent. Furthermore, if we change the initial condition so that $u(x, 0) \sim \mathrm{e}^{-ax}$, as $x \to \infty$, we find that $c = 1/a + a$, for $a > 1$, as expected [1].

#### 1.2 Fisher-Stefan equation

To obtain numerical solutions of the Fisher-Stefan problem,

$$\frac{\partial u}{\partial t} = \frac{\partial^2 u}{\partial x^2} + u(1-u), \tag{5}$$

for $0 < x < L(t)$ and $t > 0$, we first use a boundary fixing transformation $\xi = x/L(t)$ [2] so that we have

$$\frac{\partial u}{\partial t} = \frac{1}{L^2(t)}\frac{\partial^2 u}{\partial \xi^2} + \frac{\xi}{L(t)}\frac{\mathrm{d}L(t)}{\mathrm{d}t}\frac{\partial u}{\partial \xi} + u(1-u), \tag{6}$$

on the fixed domain $0 < \xi < 1$ and $t > 0$. Here $L(t)$ is the length of the domain that we will discuss later. To close the problem we must also transform the appropriate boundary conditions giving

$$\frac{\partial u}{\partial \xi} = 0 \quad \text{at} \quad \xi = 0, \tag{7}$$

$$u = 0 \quad \text{at} \quad \xi = 1. \tag{8}$$

We spatially discretise Equations (6)-(8) with a uniform finite difference mesh, with spacing $\Delta\xi$, approximating the spatial derivatives using a central finite difference approximation, giving

$$\begin{aligned}\frac{u_i^{j+1} - u_i^j}{\Delta t} &= \frac{1}{(L^j)^2}\left(\frac{u_{i-1}^{j+1} - 2u_i^{j+1} + u_{i+1}^{j+1}}{\Delta\xi^2}\right) \\ &+ \frac{\xi}{L^j}\left(\frac{L^{j+1} - L^j}{\Delta t}\right)\left(\frac{u_{i+1}^{j+1} - u_{i-1}^{j+1}}{2\Delta\xi}\right) + u_i^{j+1}(1 - u_i^{j+1}),\end{aligned} \tag{9}$$

for $i = 2, \ldots, m-1$, where $m = 1/\Delta\xi + 1$ is the total number of spatial nodes on the finite difference mesh, and the index $j$ represents the time index so that $u_i^j \approx u(\xi, t)$, where $\xi = (i-1)\,\Delta\,\xi$ and $t = j\Delta t$.

Discretising Equations (7)-(8) leads to

$$u_2^{j+1} - u_1^{j+1} = 0, \tag{10}$$

$$u_m^{j+1} = 0. \tag{11}$$

We use Newton-Raphson iteration to solve the non-linear system defined by Equations (9)-(11) and we continue with the iterations until the infinity norm of the difference between successive estimates of $u_i^{j+1}$ falls below some small tolerance, $\epsilon$. As the Newton-Raphson iterates converge we also update the $L(t)$ by considering the Stefan boundary condition

$$\frac{\mathrm{d}L(t)}{\mathrm{d}t} = -\kappa \frac{\partial u}{\partial x}, \quad \text{at} \quad x = L(t). \tag{12}$$

To incorporate the Stefan boundary condition into our numerical method we must transform the boundary condition to the fixed domain,

$$\frac{\mathrm{d}L(t)}{\mathrm{d}t} = -\frac{\kappa}{L(t)} \frac{\partial u}{\partial \xi}, \quad \text{at} \quad \xi = 1, \tag{13}$$

and we then discretise Equation (13) allowing us to update $L(t + \Delta t)$ as the Newton-Raphson iterates converge

$$L^{j+1} = L^j - \frac{\Delta t \kappa}{L^j} \left( \frac{u_m^{j+1} - u_{m-1}^{j+1}}{\Delta \xi} \right). \tag{14}$$

To demonstrate the accuracy of our numerical method to solve the Fisher-Stefan problem we consider a closely related, but simplified problem, that has an exact solution [3]. We consider

$$\frac{\partial u}{\partial t} = \frac{\partial^2 u}{\partial x^2}, \tag{15}$$

on $0 < x < L(t)$ for $t > 0$, with a moving boundary at $L(t)$. The boundary conditions are given by

$$\frac{\partial u}{\partial x} = -\mathrm{e}^t \quad \text{at} \quad x = 0, \tag{16}$$

$$u = 0 \quad \text{at} \quad x = L(t), \tag{17}$$

with the Stefan condition is

$$\frac{\mathrm{d}L(t)}{\mathrm{d}t} = -\frac{\partial u}{\partial x} \quad \text{at} \quad x = L(t), \tag{18}$$

With $L(0) = 0$, the exact solution to this moving boundary problem is

$$u(x,t) = -\mathrm{e}^{(t-x)} - 1, \tag{19}$$

on $0 < x < t$, for $0 < t < 1$. Therefore, by setting $\tilde{\lambda} = 0$, $\kappa = 1$ and changing the boundary condition from $\partial u/\partial x = 0$ at $x = 0$ to $\partial u/\partial x = -\mathrm{e}^t$ at $x = 0$, our numerical scheme ought to approximate the exact solution, Equation (19). Since the initial condition for the exact solution has $L(0) = 0$, we make progress by evaluating the exact solution at $t = \tau < 1$ and we use this solution as the initial condition from which the numerical solution can be evaluated for $t > \tau$. Results in Figure 2 compare numerical solutions and this exact solution with $\tau = 0.2$. This exercise gives us confidence in our numerical solution of the moving boundary problem since the numerical and exact solutions in Figure 2 are indistinguishable at this scale.

#### 1.3 Numerical estimate of $c$

In this work we solve both the Fisher-KPP and the Fisher-Stefan models and use the time-dependent solutions to provide an estimate of the travelling wave speed, $c$. To obtain this estimate we specify a contour value, $u(x,t) = u^*$. At the end of each time step in we use linear interpolation to estimate $x^*$ such that $u(x^*,t) = u^*$. Therefore, at the end of each time step we have estimates of both $x^*(t + \Delta t)$ and $x^*(t)$, allowing us to estimate the speed at which the contour moves

$$c = \frac{x^*(t + \Delta t) - x^*(t)}{\Delta t}. \tag{20}$$

We find that evaluating Equation (20) at each time step leads to a time series of estimates of $c$, and we find that these estimates asymptote to some positive constant value as $t \to \infty$ for those problems that support a travelling wave solution. For all results presented we set $u^* = 0.5$, but we find that our estimates of $c$ obtained using this approach are not particularly sensitive to our choice of $u^*$ [4].

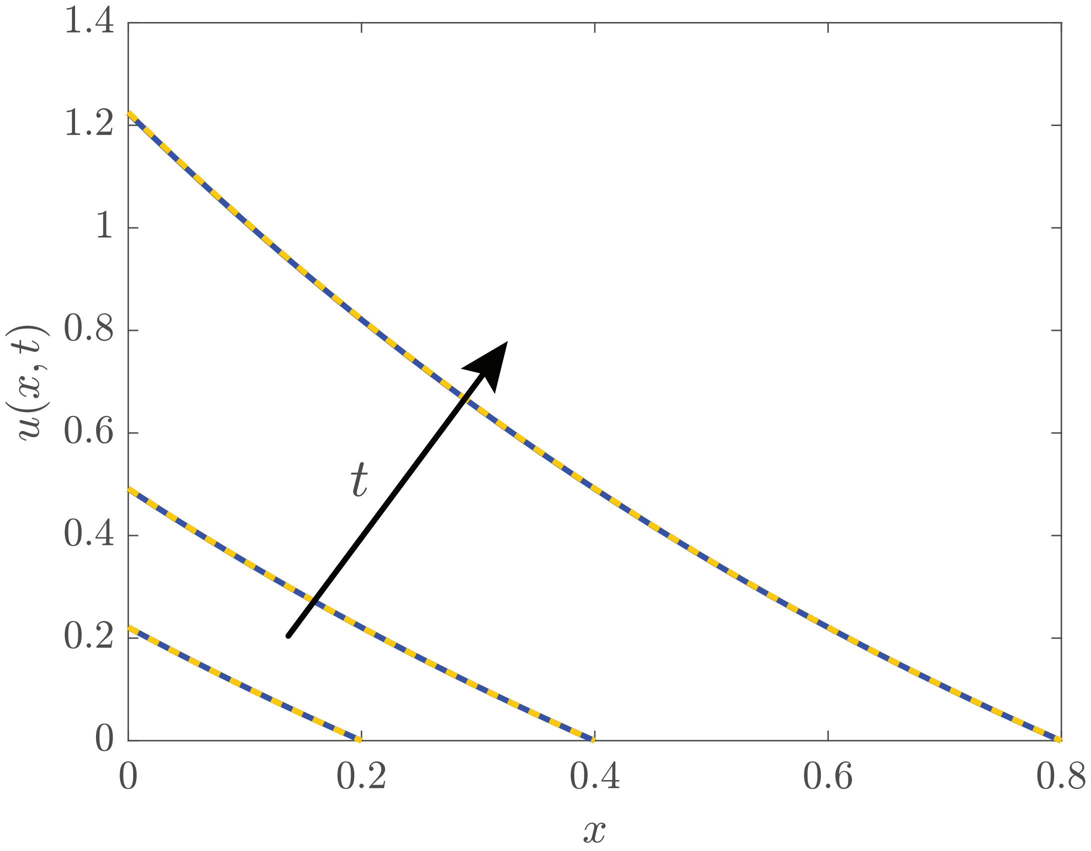


Figure 2: **Comparison of numerical and exact solutions for the simplified moving boundary problem**. Numerical solutions are obtained with $\Delta\xi = 1 \times 10^{-4}$, $\Delta t = 1 \times 10^{-3}$ and $\epsilon = 1 \times 10^{-8}$. The numerical solutions (blue solid) are superimposed on the exact solutions (yellow dashed) and solutions are shown at $t = 0.2, 0.4$ and $0.8$, with the arrow showing the direction of increasing $t$.

##### 1.4 Construction of the phase planes

The dynamical system that defines the phase plane for travelling wave solutions of the Fisher-KPP and Fisher-Stefan models is given by

$$\frac{\mathrm{d}U}{\mathrm{d}z} = V, \tag{21}$$

$$\frac{\mathrm{d}V}{\mathrm{d}z} = -cV - U(1-U). \tag{22}$$

Using the chain rule, Equations (21)-(22) can be written equivalently as

$$\frac{\mathrm{d}V}{\mathrm{d}U} = \frac{-cV - U(1-U)}{\mathrm{d}V}, \tag{23}$$

where $V = V(U)$.

When we construct phase planes in the main document we use a combination of exact and computational techniques. The locations of equilibrium points and boundary conditions are plotted on the phase plane using exact mathematical expressions for their location. The flow, defined exactly by Equations (21)-(22), is plotted using the `quiver` function in MATLAB [7]. To estimate the trajectories in the phase plane we integrate Equations (21)-(22) numerically using the `ODE45` function in MATLAB [6]. When we compute the phase plane trajectories we set the tolerance to $1 \times 10^{-4}$ in `ODE45` and we choose the initial condition using information from the linear analysis nearby the $(1, 0)$ equilibrium point to ensure that the initial condition is close to the heteroclinic orbit.

### References

[1] Murray JD. 2002. *Mathematical biology I. An introduction* New York: Springer.

[2] Simpson MJ 2015. Exact solutions of linear reaction-diffusion processes on a uniformly growing domain: criteria for successful colonization. *PLoS One* **10**, e0117949. (doi:10.1371/journal.pone.0117949).

[3] Kutluay S, Bahadir AR, Özdeş A. 1997. The numerical solution of one-phase classical Stefan problem. *J. Comput. Appl. Math.* **81**, 135-144. (doi:10.1016/S0377-0427(97)00034-4).

[4] Landman KA, Simpson MJ, Slater JA, Newgreen DF. 2005. Diffusive and chemotactic cellular migration: smooth and discontinuous travelling wave solutions. *SIAM J. Appl. Math.* **65**, 1420–1442. (doi:10.1137/040604066).

[5] Simpson MJ, Landman KA. 2006. Characterizing and minimizing the operator split error for Fisher's equation. *Appl. Math. Lett.* **19**, 604–612. (doi:10.1016/j.aml.2005.08.011).

[6] MathWorks `ode45`. Retrieved from http://mathworks.com/help/matlab/ref/ode45.html in June 2019.

[7] MathWorks `quiver`. Retrieved from http://au.mathworks.com/help/matlab/ref/quiver.html in June 2019.
